# Conserved nucleocytoplasmic density homeostasis drives cellular organization across eukaryotes

**DOI:** 10.1101/2023.09.05.556409

**Authors:** Abin Biswas, Omar Muñoz, Kyoohyun Kim, Carsten Hoege, Benjamin M. Lorton, David Shechter, Jochen Guck, Vasily Zaburdaev, Simone Reber

**Affiliations:** Max Planck Institute for Infection Biology, 10117 Berlin, Germany; Max Planck Institute for the Science of Light, 91058 Erlangen, Germany; Max-Planck-Zentrum für Physik und Medizin, 91058 Erlangen, Germany; Friedrich-Alexander-Universität Erlangen-Nürnberg (FAU), 91058 Erlangen, Germany; Max Planck Institute of Molecular Cell Biology & Genetics, 01307 Dresden, Germany; Albert Einstein College of Medicine, Bronx, NY 10461, USA; University of Applied Sciences Berlin, 13353 Berlin, Germany

**Author notes:** These authors contributed equally to this work (Abin Biswas, Omar Muñoz, Kyoohyun Kim).

## Abstract

The packing and confinement of macromolecules in the cytoplasm and nucleoplasm has profound implications for cellular biochemistry. How intracellular density distributions vary and affect cellular physiology remains largely unknown. Here, we show that the nucleus is less dense than the cytoplasm and that living systems establish and maintain a constant density ratio between these compartments. Using label-free biophotonics and theory, we show that nuclear density is set by a pressure balance across the nuclear envelope *in vitro*, *in vivo* and during early development. Nuclear transport establishes a specific nuclear proteome that exerts a colloid osmotic pressure, which, assisted by entropic chromatin pressure, draws water into the nucleus. Using *C. elegans*, we show that while nuclear-to-cytoplasmic (N/C) *volume* ratios change during early development, the N/C *density* ratio is robustly maintained. We propose that the maintenance of a constant N/C *density* ratio is the biophysical driver of one of the oldest tenets of cell biology: the N/C *volume* ratio. In summary, this study reveals a previously unidentified homeostatic coupling of macromolecular densities that drives cellular organization with implications for pathophysiologies such as senescence and cancer.

## Introduction

The cytoplasm and nucleoplasm of eukaryotic cells are complex aqueous solutions of macromolecules, small organic molecules, and ions. Both the cytoplasm and the nucleoplasm are active, self-organizing systems that exhibit cell-type and cell-cycle specific emergent properties (1,2). One fundamental and defining material property of a cell’s interior is density (dry mass per unit of volume). The packing of macromolecular components in the cytoplasm and the nucleoplasm generates environments with specific physical properties that can affect diffusion, macromolecular crowding, and enzymatic reaction rates (3–8). For a given cell type, cytoplasmic density is tightly controlled and loss of cell density homeostasis correlates with altered cell function and disease states (9–11). A dilute cytoplasm, for example, was shown to impair gene expression, cell cycle progression, and cell signalling contributing to loss of cell function during senescence (12). So, while it is evident that overall changes in intracellular density have far-reaching consequences for cellular physiology, density distributions at the subcellular level remain largely unexplored. For example, the nucleus is usually considered as a strongly crowded environment (13, 14), which harbors millions of protein complexes in addition to the genome. Recent experimental evidence, however, shows that nuclei have a lower refractive index (RI) when compared to the cytoplasm (15). Cells keep the lower nuclear RI throughout the cell cycle, even when physically and chemically challenged (16). This hints towards a lower nuclear density that is robustly maintained by the cell. Yet, in some specialized cells or growth conditions nuclei have been reported to be denser than the cytoplasm (17, 18). Thus, it remains unknown how intracellular densities are established and maintained across cell compartments.

Just like cellular density, cell size is narrowly constrained within a specific cell type (19–21). Although it has recently been observed that cytoplasmic density decreases when cells grow too large (12), it remains unknown whether and how cell density and volume are mechanistically linked. Cell size, however, is one of the critical parameters controlling the size of intracellular structures (22–24). A well-known example is the constant nuclear-to-cytoplasmic (N/C) volume ratio first described by Richard Hertwig as early as 1903 (25). Despite a century of size-scaling observations, the mechanisms coordinating the growth and size of subcellular structures remain largely elusive.

In this study, we show that nuclei are consistently less dense than their surrounding cytoplasm. We find a homeostatic coupling of the nuclear volume to the cytoplasm by a pressure balance mechanism across the nuclear envelope that maintains a constant density ratio of 0.8 between these two compartments. While absolute densities significantly differ between species, we discover the nuclear-to-cytoplasmic *density* ratio to be conserved in 10 model systems across the eukaryotic kingdom, suggesting that this measure is a fundamental cellular characteristic. To mechanistically decipher how nuclear density is established, we reconstitute nuclear assembly and growth in *Xenopus* egg extracts. We find that nuclear protein complexes establish a colloid osmotic pressure, which — assisted by entropic chromatin pressure — inflates the *Xenopus* nucleus to its observed volume and results in a nuclear density that is lower than that of the cytoplasm. We substantiate these *in vitro* findings by observations in early *C. elegans* embryos, where N/C *density* ratios are robustly maintained throughout early development even when N/C *volume* ratios change. Based on general biophysical principles of pressure balance and kinetics of active transport, we present a unifying theoretical framework that robustly predicts nuclear density and explains how density responds to environmental changes. In summary, we reveal a previously unidentified subcellular regulation of macromolecular crowding, which establishes intracellular pressures driving cellular organization.

## Results

### During assembly, the *Xenopus* nucleus establishes a lower density than the surrounding cytoplasm

While it is known that overall cell density shows little variation within a given cell type (10, 26), there is limited understanding on how density varies at the subcellular scale. To close this gap in knowledge, we used correlative fluorescence and optical diffraction tomography (ODT, Figure S1A, 27, 28) to identify subcellular structures using fluorescence and measure their corresponding 3D refractive index in individual cells (Figure 1A). In biological materials, the RI is linearly proportional to the dry mass density (ρ) of constituent macromolecules (28–32). Thus, these RI tomograms provide a direct, label-free, and quantitative readout of cellular mass and density distributions (Materials and Methods). In a set of 10 model systems, ranging from yeast to human cells, we consistently found the nucleoplasm to be less dense than the cytoplasm (Figure 1B). While the absolute cytoplasmic and nucleoplasmic densities differed between species (Figure S1B), the ratio of nuclear-to-cytoplasmic densities was invariantly 0.78 ± 0.01 (arithmetic mean ± SEM, Figure 1C), suggesting this ratio is actively maintained by the cell. These observations imply that after mitosis, when the nucleoplasm and cytoplasm are fully mixed, the assembling nucleus needs to reestablish a lower density. The formation of eukaryotic nuclei upon mitotic exit therefore provides an experimental system to examine the mechanisms that control nuclear density. To this aim, we reconstituted nuclear assembly *in vitro* using *X. laevis* egg extracts. Such nuclei are known to recapitulate many essential processes such as DNA replication, nucleocytoplasmic transport, and nuclear growth (33, 34). We visualized chromatin (Figure 1D, DNA, Figure S1C), membranes (Figure S1D), chromatin replication (Figure S1C) and nuclear volume over time (Figure 1E). Additionally, we visualized and quantified nuclear import by the accumulation of green fluorescent protein (GFP) fused to a nuclear localization signal (NLS) (Figure 1D, NLS-GFP, 1F). Simultaneously, we measured the RI tomograms and calculated nuclear dry mass, which gradually increased over time (Figure 1D, RI, 1E). At 60 minutes, the average reconstituted *X. laevis* nucleus had a dry mass of 51 ±2 pg. Combining volumetric and dry mass information allowed us to calculate the density of assembling and growing nuclei over time (Figure 1G, S1E).

**Figure 1.**
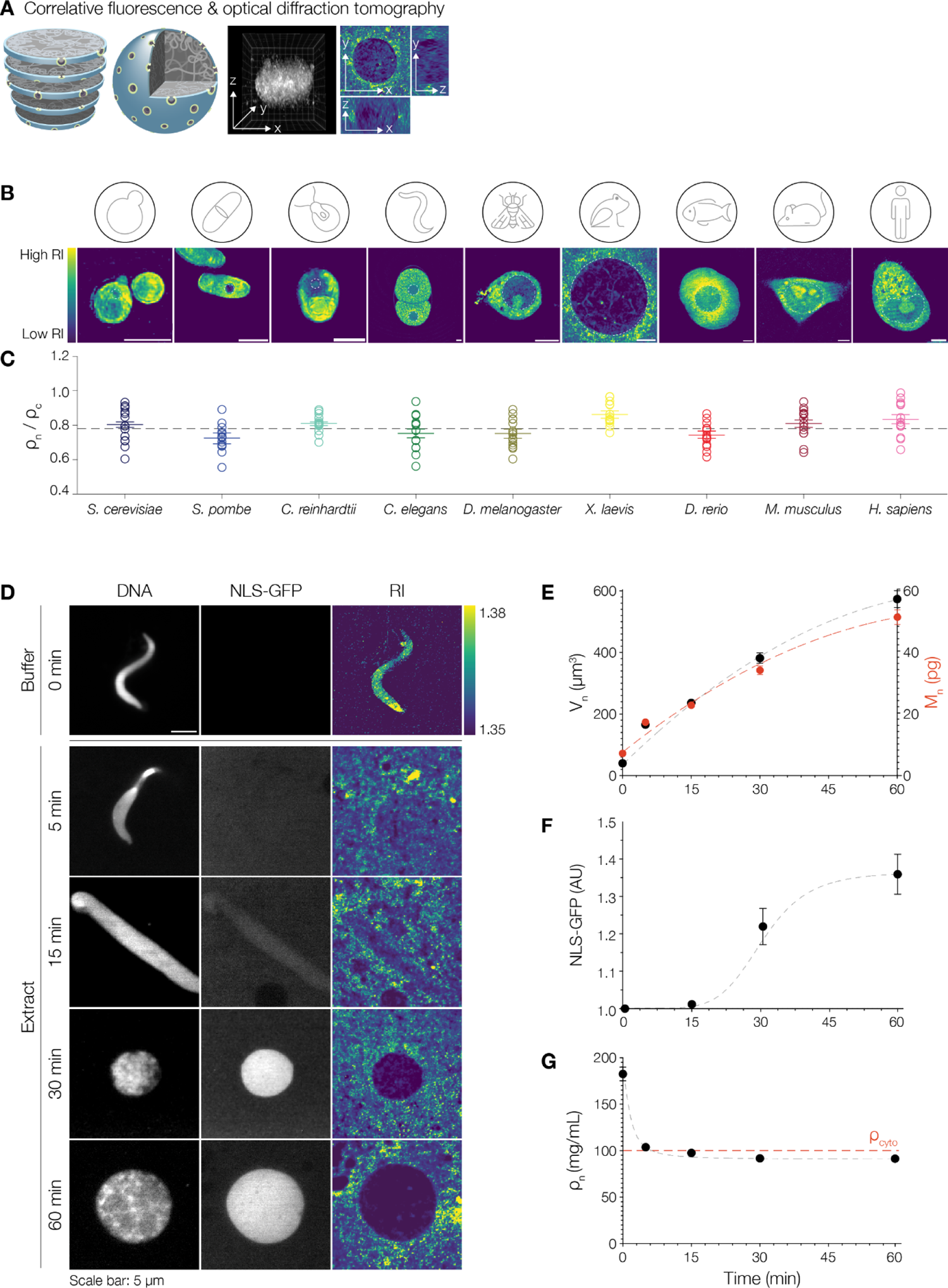
During assembly, the *Xenopus* nucleus establishes a lower density than the surrounding cytoplasm. (A) A correlative confocal fluorescence and optical diffraction tomography (ODT) setup was used to obtain 3D volume and refractive index (RI) distributions of nuclei (schematic). Middle panel: Volumetric view of a representative nucleus where DNA was stained with Hoechst-33342. Right panel: xy, yz and xz projections of a nucleus showing the RI distributions, as obtained by ODT. See also Figure S1A. (B) Representative images of cells from different eukaryotes imaged via ODT. Scale bar: 5 μm. Colors in the images show the RI distribution from low (dark blue) to high (yellow). See also Figure S1B for absolute values of nuclear and cytoplasmic mass densities. (C) Ratio of nuclear and cytoplasmic density (ρ) across different eukaryotes. Each circle represents the ratio of the N/C density of one cell. Nuclear densities were measured in nucleoplasmic regions excluding nucleoli. 15 cells were analyzed per organism. Bars show mean ± SEM. The average ratio of the N/C density is 0.78 ± 0.01 (indicated by dashed line). (D) Representative images at different time points of nuclear assembly. Demembranated sperm nuclei were added to buffer (0 min) or to *Xenopus* egg extracts (5 - 60 min). Left panel: Fluorescence images of DNA stained with Hoechst-33342. Middle panel: Fluorescence images of NLS-GFP. Scale bar: 5 μm. Right panel: Central slice from a reconstructed ODT tomogram. Bar on the right shows the RI distribution range. (E) As nuclei assemble, nuclear volume and dry mass increases. Volume (V_n_, mean ± SEM) at indicated time points: V_n_0_ = 40 ± 2 μm^3^ (n = 30), V_n_5_= 171 ± 6 μm^3^ (n = 160), V_n_15_= 236 ± 7 μm^3^ (n = 170), V_n_30_ = 382 ± 17 μm^3^ (n = 173), and V_n_60_ = 573 ± 27 μm^3^(n = 175). Dry mass (M_n_, mean ± SEM) at indicated time points: M_n_0_ = 7.2 ± 0.4 pg (n = 30), M_n_5_ = 17.3 ± 0.5 pg (n = 160), M_n_15_ = 22.8 ± 0.6 pg (n = 170), M_n_30_ = 34.0 ± 1.4 pg (n = 173) and M_n_60_ = 51.2 ± 2.4 pg (n = 175). In panels (E-G), dashed lines are used to guide readers along the trend followed by the experimental data. (F) Nuclear GFP-NLS fluorescence intensity (in arbitrary units, AU) was measured at each time point (n = 18 from 2 independent experiments). (G) The average nuclear mass density (ρ_n_) was calculated at different time points. As nuclei assemble, the mass density drops from p_n_0min_ = 183 ± 7 mg/mL (n = 30) to p_n_5min_ = 104 ± 1 mg/mL (n = 160). Between 5-15 minutes, the nuclear mass density reduces below that of the cytoplasm (p_Gyto_ = 100 ± 2 mg/mL, red dashed line) to p_n_15min_ = 97.6 ± 0.9 mg/mL (n = 170), p_n_30min_ = 91.7 ± 0.9 (n = 173) and p_n_60min_ = 91.4 ± 0.8 (n = 175).

Once densely packed sperm chromatin (p_n_0min_ = 183 ± 7 mg/mL, Figure 1D and G, 0 min) was added to *Xenopus* egg extract, its density quickly decreased within five minutes (p_n_5min_ = 104 ± 1 mg/mL) to level with that of the cytoplasm (ρ_cyto_ = 100 ± 2 mg/mL, Figure 1D and G, 5 min). Concurrent with the start of nuclear import, nuclear density further reduced and then steadily remained below the density of the cytoplasm while the nuclei continued to grow (Figure 1D and G, p_n_15min_ = 97.6 ± 0.9 mg/mL, p_n_30min_ = 91.7 ± 0.9 mg/mL, p_n_60min_ = 91.4 ± 0.8 mg/mL). Thus, our correlative fluorescence and ODT set-up allowed us to systematically measure cytoplasmic and nuclear densities with high spatial (120 nm) and temporal (1 sec) resolution (Figure S1F and G). We found that the *Xenopus* nucleus robustly establishes a lower density than the surrounding cytoplasm within 15 minutes of nuclear assembly. Such a constant N/C density ratio of approximately 0.8 — ubiquitous to nine other species across the eukaryotic kingdom — suggests that this measure is a fundamental cellular characteristic. We next ventured to identify the key biochemical and biophysical mechanisms responsible for the observed dynamics in nuclear density.

### Nucleoplasmin-dependent chromatin decondensation and nuclear import lower nuclear density

The decrease in nuclear density as described in Figure 1 can be nominally divided into two phases. A first fast decrease in density that happens concomitantly with apparent sperm chromatin decondensation. This is followed by a second — more gradual — decrease in density which starts concurrently with nucleocytoplasmic transport and results in a nuclear density below that of the cytoplasm (Figure 2A). What are the molecular processes responsible for the decrease of nuclear density? It is known that highly compact *Xenopus* sperm chromatin undergoes rapid Nucleoplasmin-dependent decondensation in *Xenopus* egg extracts. Nucleoplasmin (Npm2) is a pentameric embryonic histone chaperone that removes sperm-specific basic proteins, binds core histones, and promotes their assembly into nucleosomes (35–39). To quantify how chromatin decondensation affects its density, we supplied *Xenopus* sperm chromatin (in buffer) with purified recombinant Npm2 (Figure S2A) and measured chromatin density over time: within 10 minutes, chromatin rapidly decondensed (Figure 2B) and increased in volume (Figure S2B) while its density decreased (Figure 2C). Next, to show that Npm2 is both necessary and sufficient for the initial reduction in nuclear density, we performed immunodepletion and add-back experiments (Figure 2D). Consistent with previous reports (35, 36), Npm2 depletion left the sperm chromatin condensed, and adding back Npm2 led to its quick decondensation (Figure 2E, Figure S2C-E). In Npm2-depleted extracts, sperm chromatin had a higher density than in control-depleted extracts (Figure 2F, p_10min_ = 115 ± 5 mg/mL versus 98 ± 5 mg/mL), suggesting that Npm2-dependent decondensation is necessary for the initial drop in density. Sperm nuclei in control-depleted and Npm2-supplied extracts had a similar density as control nuclei (Figure 2F, p_10min_ = 98 ± 5 mg/mL versus 89 ± 3 mg/mL), suggesting that Npm2-dependent chromatin decondensation is sufficient for the initial reduction in nuclear density.

**Figure 2.**
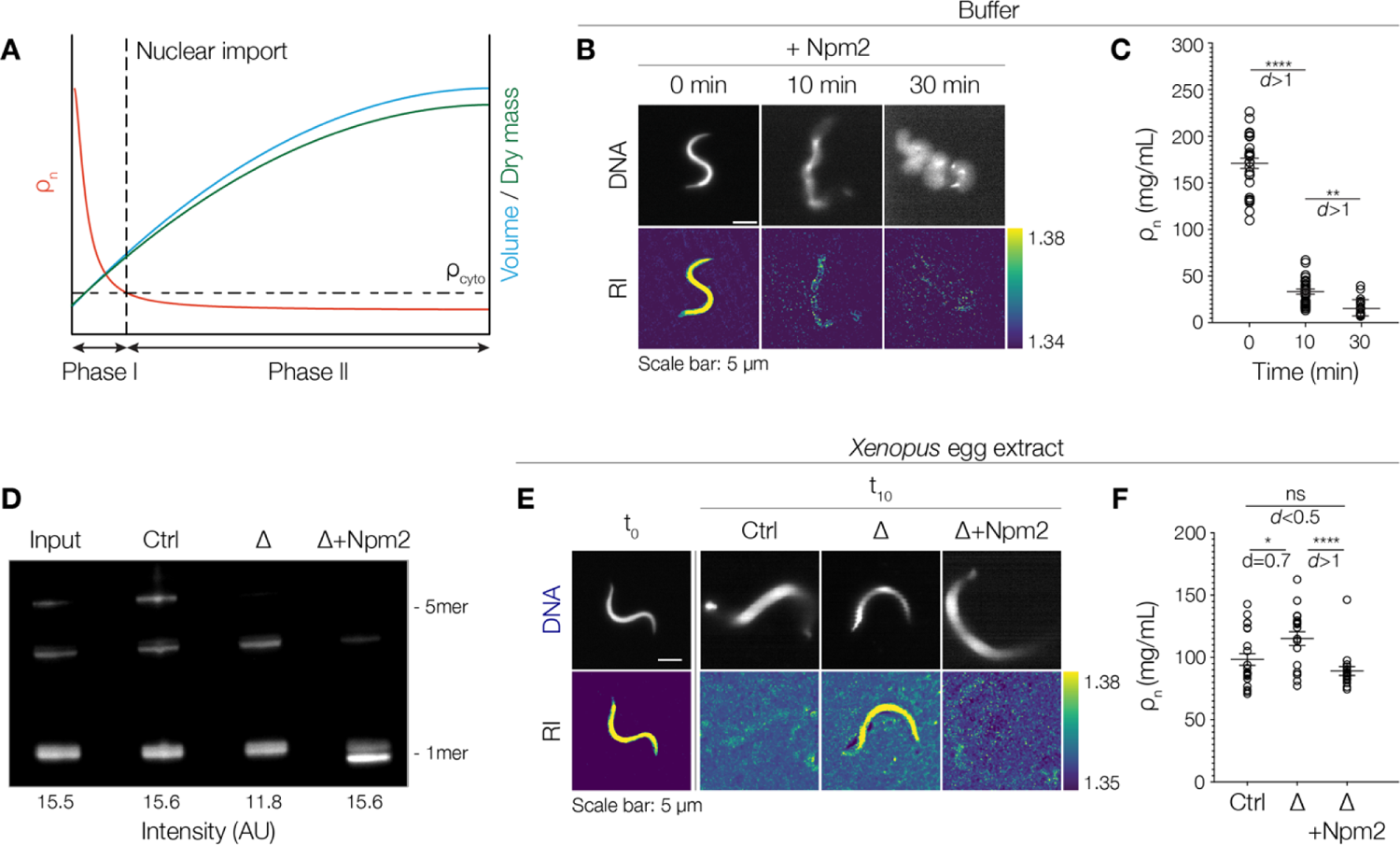
Nucleoplasmin is both necessary and sufficient to reduce nuclear density to that of its colvent. (**A**) Schematic summarizing changes in nuclear density (p_n_, red), volume (blue), and dry mass (green) during nuclear assembly. Final nuclear density is lower than that of the cytoplasm (pc_yto_). (B) Adding purified Npm2 (final concentration: 9 pM) to sperm nuclei in buffer reduces the RI and increases volume (Figure S2B). Representative images at different time points. Top panel shows DNA stained with Hoechst-33342. Scale bar: 5 pm. Lower panel shows the central slice from the reconstructed ODT tomogram (RI). Bar on the right shows the RI distribution range. See also Figure S2. (**C**) Quantification of chromatin density (pn) at 10 and 30 minutes after the addition of purified Npm2. Circles represent individual data points from 3 independent experiments, n = 30, 27, and 23 at 0, 10, and 30 min, respectively. Bars indicate the mean ± SEM in all graphs. p_n_0min_ = 171.0 ± 5.5 mg/mL, p_n_10min_ = 33.4 ± 2.8 mg/mL, p_n_30min_ = 15.3 ± 2.1 mg/mL. Mann-Whitney test where ** indicates p < 0.01 and **** indicates p < 0.0001. Cohen’s *d*_0_&_10min_> 1, Cohen’s *d* _10_&_30min_> 1. (D) Immunoblot of *Xenopus* egg extracts probed with Npm2 antibody. Input, control-depleted, Npm2-depleted (△), and Npm2-depleted with recombinant Npm2 added back to 4 pm final concentration (A+Npm2). Note that mostly the pentameric Npm2 was depleted. This, however, was sufficient to inhibit chromatin decondensation (see Figure S2E). Quantification of total intensity of all bands in arbitrary units (AU). (E) Nucleoplasmin is both necessary and sufficient to reduce the density of sperm chromatin. Representative fluorescence (top panel, DNA stained with Hoechst-33342) and RI images (bottom panel, central slice of tomogram) of assembling nuclei from control-depleted, A and A+Npm2 extracts. The t_0_ timepoint shows sperm nuclei in buffer. After 10 minutes (t_10_), sperm nuclei decondensed in control-depleted extracts but not in ANpm2 extracts. The add back of recombinant Npm2 (A+Npm2) is sufficient to decondense sperm nuclei. Scale bar: 5 pm. Bar on the right shows the RI distribution range. (F) Quantification of chromatin density (p_n_) 10 minutes after the start of nuclear assembly in control-depleted, Npm2-depleted, and Npm2-supplied extracts. p_n_ANpm2_ = 115.1 ± 5.5 mg/mL is significantly higher than p_n_MOcK_ = 98.4 ± 4.7 mg/mL and p_n_A+Npm2_ = 89.2 ± 3.6 mg/mL, n = 19, 22, and 18 samples from 2 independent experiments. Bars show mean ± SEM. Mann-Whitney where ns indicates p > 0.05, * indicates p < 0.05 and **** indicates p < 0.0001. Cohen’s *d*ct_rl_&_ANpm2_ = 0.7, Cohen’s *d*_ANpm2&+Npm2_ > ^1^ and ^Cohens^ *d*_Ctrl&A+Npm2_ < 0.5.

The further reduction in nuclear density below the density of the cytoplasm started concurrently with the accumulation of nuclear NLS-GFP suggesting that nucleocytoplasmic transport might play a role. Therefore, we assembled nuclei in the presence of import inhibitors (Figure 3A, 40, 41). The successful inhibition of nuclear import was visualized by a significant reduction of NLS-GFP accumulation in the nucleus (Figure 3A, B). Import-deficient nuclei assembled after chromatin addition and were initially similar in size. After 60 minutes, however, the import-deficient nuclei were significantly smaller when compared to control nuclei (Figure 3C, Supplemental theory, Figure S9) consistent with previous reports (42–44). Importantly, the density of the import-deficient nuclei (p_n__60m_in_ = 101.1 ±1.6 mg/mL) did not reduce below the density of the cytoplasm implying that nuclear import is essential to further reduce nuclear density (Figure 3A, D, E; Supplemental theory, Figure S9).

**Figure 3.**
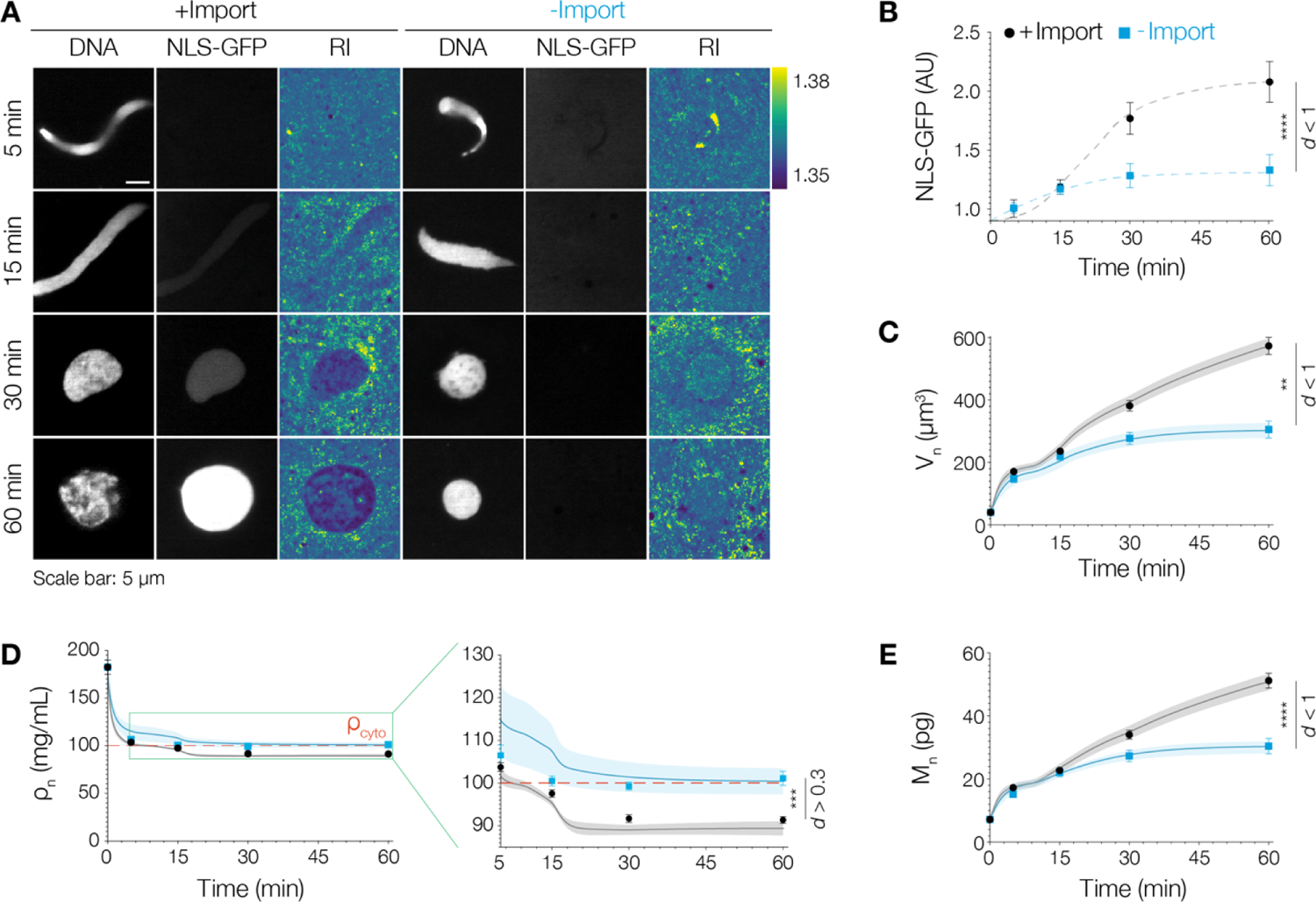
Nuclear import is required to reduce nuclear density below that of the cytoplasm. (A) Nuclei were assembled in *Xenopus* egg extract in the absence (+Import) or presence of import inhibitors (-Import). Left panels: fluorescence images of DNA stained with Hoechst-33342, middle panel: fluorescence images of NLS-GFP and right panel: central slice from the reconstructed ODT tomogram. Scale bar: 5 μm. Bar on the right shows the RI distribution range. (B) Quantification of NLS-GFP import. Nuclei assembled in the presence of inhibitors (-Import, blue squares show mean at different time points) accumulated significantly lower NLS-GFP protein in comparison to control nuclei (+Import, black circles show mean at different time points). n = 30 from 3 independent experiments. Bars indicate SEM in all graphs. Mann-Whitney test where **** indicates p < 0.0001. Cohen’s *d*_+Impor_t&-_Import_ < 1. Dashed lines are used to guide readers along the trend followed by the experimental data. (C) Quantification of nuclear volume. Volume (blue squares, mean ± SEM) at indicated time points for import deficient nuclei V_n_5_ = 148 ±123 μm^3^(n = 30), V_n_15_= 221 ± 14 μm^3^(n = 30), V_n_30_ = 278 ± 19 μm^3^ (n = 30) and V_n_60_ = 306 ± 27 μm^3^ (n = 25). Black circles show mean at different time points for control nuclei. At 60 minutes, there is a significant difference between the volumes of import-deficient and control nuclei. Mann-Whitney test where ** indicates p < 0.01. Cohen’s *d*_+Impor_t&-_Import_< 1. In panels (C-E), bold lines and shaded areas represent the mean ± SEM from the theoretical simulations for the control (+Import, black) and import deficient (-Import, blue) conditions. See also Supplemental theory. (D) Quantification of nuclear density upon import inhibition. ρ_n_ (blue squares, mean ± SEM), at indicated time points for import deficient nuclei p_n_5min_ = 106.5 ± 2.4 mg/mL (n = 30), p_n_15min_= 100.5 ± 1.2 mg/mL (n = 30), p_n_30min_ = 99.3 ± 0.9 mg/mL (n = 30) and p_n_60min_= 101.1 ±1.6 mg/mL (n = 25). Import-deficient nuclei have a density, p_n_, similar to that of the cytoplasm (red line). Black circles show mean at different time points for control nuclei. Inset shows a zoomed-in view of ρ_n_ between 5-60 minutes. Mann-Whitney test where *** indicates p < 0.001. Cohen’s *d*_+Impor_t&-_Import_ > 0.3. (E) Quantification of dry mass upon import inhibition. M_n_ (blue squares, mean ± SEM) at indicated time points for import deficient nuclei M_n_5_ = 15.2 ± 1.1 pg (n = 30), M_n_15_ = 22.1 ± 1.4 pg (n = 30), M_n_30_ = 27.3 ± 1.8 pg (n = 30) and at M_n_60_ = 30.4 ± 2.4 pg (n = 25). At 60 minutes, there is a significant difference between the dry mass of import-deficient and control nuclei. Mann-Whitney test where **** indicates p < 0.0001. Cohen’s *d*_+Import_&-_Impor_t < 1.

Consistently, when we simultaneously inhibited chromatin decondensation and nuclear pore formation, nuclei retained a density as high as that of condensed sperm chromatin (Figure S3A-D). Taken together, we have established that (1) Npm2-dependent chromatin decondensation quickly equalizes nuclear density to that of the surrounding cytoplasm and that (2) nuclear import is essential to further decrease nuclear density below that of the cytoplasm. The question then arises how active import of macromolecules into the nucleus can establish a reduced nuclear density?

### Osmotically active solute macromolecules and chromatin determine nuclear volume and density

Nuclear import results in a reduction in nuclear density. This, at first glance, seems counterintuitive. How can the addition of molecules to the nucleus make it less dense? It has been proposed that the import of nuclear proteins generates an osmotic pressure across the nuclear envelope that inflates the nucleus through water influx (45–47).

This has led to the distinct prediction that at steady-state the concentration of proteins in the nucleus and the cytoplasm should be equal (48–50). Since we can directly measure densities, we estimated the protein concentration of the nucleoplasm n_n_ and the cytoplasm n_c_ to be 0.51 ± 0.02 mM and 0.65 ± 0.02 mM, respectively (Figure 4A, Supplemental theory, Table S1). Here, it is important to note that we used the average mass of protein complexes, which is 155 kDa for nuclear proteins and 131 kDa for cytoplasmic proteins (51). This significantly lowers the number estimate with consequences for the colloid osmotic pressures (cf. 49 and Supplemental theory, Table S1). To make this tangible, consider a ribosome: assembling more than 80 proteins into a single macromolecular complex reduces their osmotic pressure by a factor of 86 (46). Based on the above concentrations, we then calculated the osmotic pressure exerted by nuclear and cytoplasmic proteins (Figure 4A). The final pressure difference of ΔP = −330 Pa, however, would compress the nucleus. This means that the osmotic pressure exerted by nuclear proteins alone is not sufficient to draw enough water into the nucleus to explain the observed final nuclear volume and density.

**Figure 4.**
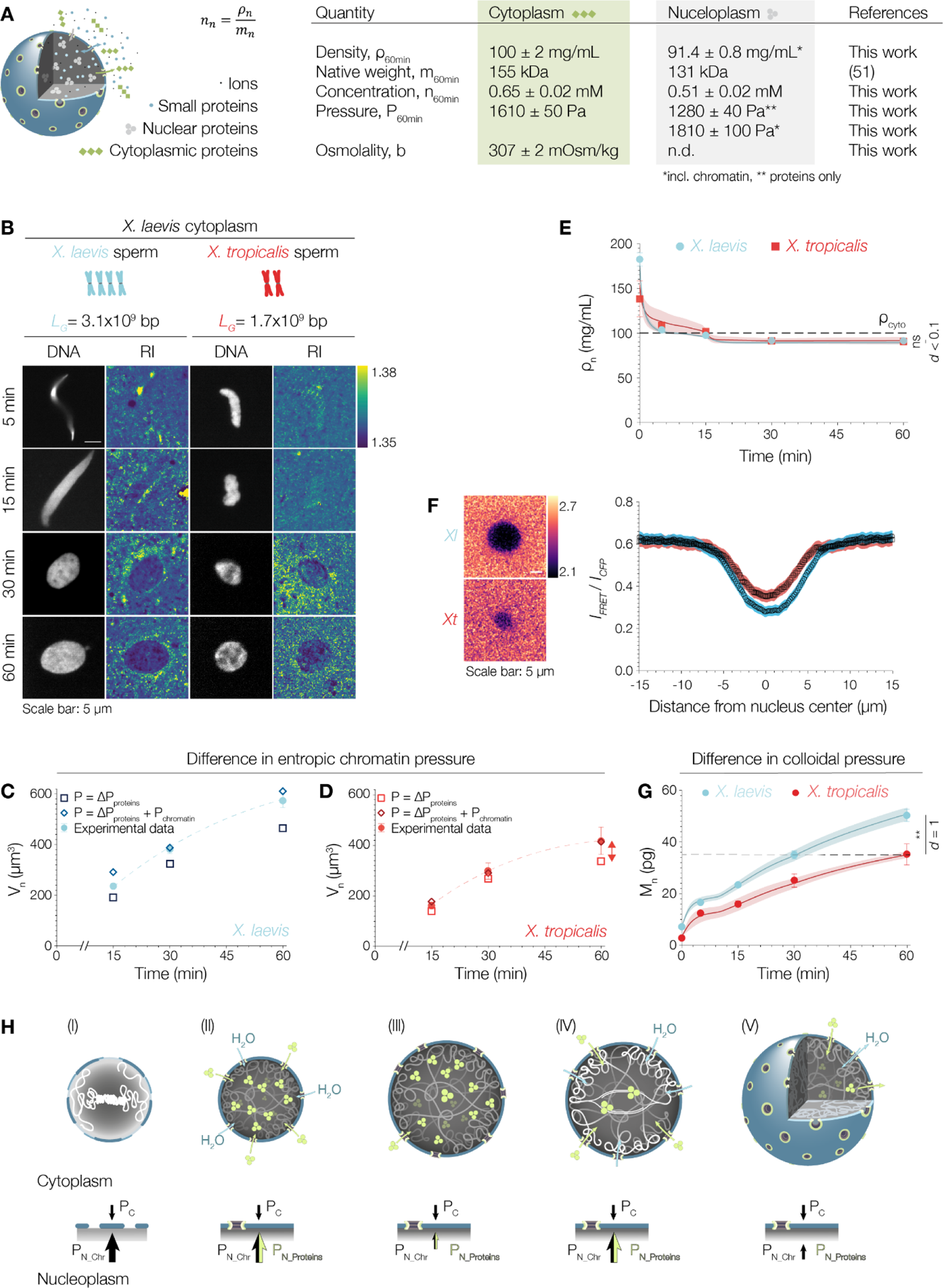
Osmotically active solute macromolecules determine nuclear density and volume. (A) Nuclear transport results in the establishment of a specific nuclear proteome that is distinct from the cytoplasmic protein composition. Small proteins (< 40 kDa), ions, and small metabolites - while being major osmolytes - can freely diffuse through nuclear pore complexes, which allows them to reach chemical equilibrium. The main contributors to the pressure difference across the nuclear envelope are proteins that preferentially locate to either the cytoplasm or the nucleoplasm. Quantities measured and calculated to derive pressure balance and predict nuclear volumes. For a detailed overview of the calculations please refer to the Supplemental theory and Table S1 therein. (B) Nuclear assembly reaction in *X. laevis* egg extract using either *X. laevis* sperm (light blue, genome size (LG) of 3.1 × 10^9^ base pairs) or *X. tropicalis* sperm (red, LG = 1.7 × 109 base pairs) as chromatin source. Fluorescence images of DNA stained with Hoechst-33342 and the corresponding RI images are shown at different time points. Scale bar: 5 μm. Colormap on the right shows the RI distribution in the images. (C) Quantification of nuclear volume (V_n_) around *X. laevis* (blue) and (**D**) *X. tropicalis* (red) sperm. Experimental data points (circles) and predicted values from pressure balance. Volume (mean ± SEM) at indicated time points for *X. tropicalis* V_n_0_ = 25 ± 3 μm^3^(n = 15), V_n_5_= 123 ± 13 μm^3^(n = 25), V_n_15_= 162 ± 14 μm^3^ (n = 29), V_n_30_ = 299 ± 32 μm^3^ (n = 29) and V_n_60_ = 412 ± 55 μm^3^ (n = 30). *X. tropicalis* nuclei are approx. 30% smaller than *X. laevis* nuclei. Considering colloid osmotic pressure by proteins (squares) alone results in an underestimation of nuclear volume. Including entropic chromatin pressure (diamonds) matches experimental values. (D) Quantification of p_n_ at different time points for nuclei assembled using *X. laevis* sperm and *X. tropicalis* sperm. *X. tropicalis*: ρ_n_ (mean ± SEM) at indicated time points p_n_0min_ = 138.3 ± 19.9 mg/mL (n = 15, sperm), p_n__gm_in_ = 107.5 ± 3.8 mg/mL (n = 25), p_n_16_m_in_= 101.7 ± 3.0 mg/mL (n = 29), p_n__30m_in_ = 91.3 ± 2.7 mg/mL (n = 29) and p_n_60min_ = 90.1 ± 2.2 mg/mL (n = 30). Density, pc_yto_, of the cytoplasm indicated by a black line. After 60 minutes, the nuclear densities are comparable and significantly below that of the cytoplasm. Cohen’s *d*Laevis&Tropcals<0.1. (E) A FRET probe was used to measure RanGTP levels in nuclei assembled from *Xenopus laevis* (*Xl*, blue) and *Xenopus tropicalis* (*Xt*, red) sperm nuclei. Representative images on the left. FRET ratio signal decreases within nuclei due to the high concentration of RanGTP in comparison to the cytoplasm. Line scan quantifications of the FRET ratio signal (Intensity_FRET_/Intensity_CFP_) from both nuclei (n = 39 for each condition). A larger indent in the line scans for *X. laevis* nuclei indicates an overall higher concentration of RanGTP. Scale bar: 5 μm. Color scale on the right shows the fluorescence intensity range in the images. (F) Quantification of nuclear dry mass at different time points of nuclear assembly. For *Xl* nuclei, dry mass values are similar to values shown in Figure 1E. For *Xt* nuclei (mean ± SEM), M_n_0_ = 2.9 ± 0.2 pg (n = 15), M_n_5_ = 12.6 ± 1.1 pg (n = 25), M_n_15_ = 15.9 ± 1.2 pg (n = 29), M_n_30_ = 26.3 ± 2.6 pg (n = 29) and M_n_60_ = 36.4 ± 3.9 pg (n = 30). *Xl* nuclei accumulate more dry mass than *Xt* nuclei. Dashed black line shows how the dry mass in *Xt* nuclei at 60 minutes is similar to the dry mass of *Xl* nuclei at 30 minutes. Mann-Whitney test where ** indicates p < 0.01. Cohen’s *d*_lae_v_is_&tr_ops_ = 1. Bold lines and shaded areas represent the mean ± SEM from the theoretical simulations for *X. laevis* (blue) and *X. tropicalis* (red) nuclei. See also Supplemental theory. (G) Pressure balance model. (l) Chromatin unfolds towards its thermodynamically preferred volume in the cytoplasm. (ll) At the same time, nuclear envelope assembly starts and confines chromatin to a well-defined nuclear compartment, which creates an outward pressure. Then, nucleocytoplasmic transport, with rates dependent on chromatin content, establishes nuclear identity with a net accumulation of nuclear proteins. As a consequence, nuclear dry mass increases. The imported nuclear protein complexes create an outward colloid osmotic pressure, which — assisted by entropic chromatin pressure — drives water influx into the nucleus and thus nuclear growth (lll). (lV) Replicating chromatin contributes to pressure balance both directly as a confined polymer-like chain and indirectly as a regulator of nuclear import, driving nuclear growth further (V).

What are we missing? The most evident nuclear macromolecule that has the potential to exert additional pressure is chromatin (48, 49, 52). To experimentally assess the contribution of chromatin, we assembled nuclei around either tetraploid *X. laevis* (genome size: 3.1 × 10^9^ bp) or diploid *X. tropicalis* (genome size: 1.7 × 10^9^ bp) sperm in an identical cytoplasm (Figure 4B). As reported before (42, 53), the *X. tropicalis* nuclei were smaller than *X. laevis* nuclei but importantly they had the same nuclear density (Figure 4C-E, Figure S4A). As the extract system is transcriptionally and translationally inactive and the total NPC number is similar in *X. laevis* and *X. tropicalis* nuclei (42), we propose that chromatin has a non-negligible effect on the pressure balance and thus nuclear size.

In fact, chromatin can contribute to the pressure balance both directly as a confined polymer-like chain (52) and indirectly as a regulator of nuclear import via the RanGTP gradient (54). Chromatin exerts an outward entropic pressure when the nuclear envelope confines its thermodynamically favored volume. Indeed, when we included chromatin entropic pressure to the pressure balance (Supplemental theory, Equation S5), nuclear volume predictions quantitatively matched the measured experimental values (Figure 4C, D). Thus, we estimated chromatin entropic pressure to contribute to about 20% of the final size of *Xenopus* nuclei.

Next, to quantify the effect of chromatin content on nuclear import and final mass, we measured the RanGTP gradient, NLS-GFP accumulation, and nuclear dry mass. Consistent with previous reports (55, 56), we observed a significant enrichment of RanGTP in the nucleoplasm. *X. laevis* nuclei showed higher RanGTP levels than *X. tropicalis* nuclei (Figure 4F). Corroborating this, the accumulation of NLS-GFP, as a proxy for nuclear import efficiency, was also higher for *X. laevis* nuclei (Figure S4B) leading to a 1.4-fold increase in nuclear dry mass when compared to *X. tropicalis* nuclei. Interestingly, nuclear dry mass was approximately equal at 30 min of *X. laevis* and 60 min of *X. tropicalis* nuclear assembly: this is when the nuclei are expected to have roughly the same DNA amount (tetraploid and replicated diploid, dashed line in Figure 4G). Collectively our data imply that the nuclear population of imported macromolecules creates a colloid osmotic pressure, which — assisted by entropic chromatin pressure — inflates the nucleus to its observed final volume and resulting lower density.

Despite the long-established scaling relation of nuclear size with DNA content (57, 58), recent studies moved the focus towards cytoplasmic factors setting nuclear size independent of nuclear DNA content (42, 49, 50, 59–61). As we show here, these two concepts are not mutually exclusive and can thus be linked in a unifying framework. To this end, we developed a detailed mechanistic model of nuclear transport (62, 63) that includes DNA replication and is coupled to the pressure balance allowing us to describe the full dynamics of nuclear growth (Movie S1, Supplemental theory, Equations S25 and S26). One prediction of our model is that in the absence of replication, nuclear protein concentration and therefore nuclear size would rapidly reach a steady state. Indeed, as reported previously (53), we observe that replication-deficient nuclei were significantly smaller than control nuclei and stopped growing after ∼ 30 min. As predicted (Supplemental theory, Figure S10), replication-deficient nuclei also had a significantly smaller nuclear proteome with a final dry mass of 24.2 ± 1.4 pg (Figure S4C-F). Taken together, we propose that chromatin has a direct, non-negligible effect on the pressure balance via its entropic pressure and an indirect effect by regulating nuclear import. Based on general biophysical principles of pressure balance and kinetics of active transport, our model robustly represents the volume and dry mass dynamics of nuclear assembly. It further correctly predicts the effects of biochemical perturbations, which include inhibition of nuclear import and replication, osmotic challenges (Figure S5A-E), and changes in chromatin content (Movie S1, Supplemental theory, Figures S9-S11). Next, to substantiate our mechanistic understanding, we wished to obtain independent experimental support from an *in vivo* system, in which nuclear volumes change during development.

### During early development, the nuclear-to-cytoplasmic *density* ratio is robustly maintained while *volume* ratios change

Contrary to the N/C *density* ratio, the well-known N/C *volume* ratio has been studied extensively ever since first described by Richard Hertwig as early as 1903 (25). Although N/C volume ratios are reported to be constant, they can change when cells change fate or size during development and differentiation (60, 61, 64–67). What happens to N/C density ratios when N/C volume ratios change? To study how nuclear and cytoplasmic densities respond to changing N/C volume ratios, we imaged early divisions in *C. elegans* embryos via ODT (Figure 5A, Movie S2). As cell number increased, cell volume decreased, nuclear size scaled with cell volume (Figure 5B, S6A) and N/C volume ratios changed (Figure 5C) consistent with previous reports (68–71). To iterate the role of chromatin, we in addition imaged tetraploid embryos (Figure 5A, Movie S3). Consistent with our results in *Xenopus* egg extracts, higher ploidy led to larger nuclei (Figure 5D). Interestingly, not only nuclear size but also cell and embryo size scaled with ploidy spanning four hierarchical levels of biological organization (Figure S6C-F, 72). The overall nuclear scaling behavior of diploid and tetraploid embryos, however, was comparable (Figure 5D, E, S6B). Remarkably, nuclear and cytoplasmic densities were constant over the course of the first four divisions in both worm strains (Figure 5F-I) with the nucleus always being less dense than the cytoplasm. Thus, while the N/C volume ratio increased, the N/C density ratio was robustly maintained (Figure 5J, K). Importantly, these observations are fully consistent with our pressure balance model: absolute nuclear size decreases because the maternal cytoplasm of the one-cell embryo is partitioned into smaller cells at constant density (Figure 5F, G). Simply, in smaller cells, the material available for nuclear growth is reduced. While the absolute dry mass and volume of nuclei decreases, the absolute dry mass and volume of the cytoplasm decreases even more rapidly (Figure S6G, H). Thus, to keep a constant N/C density ratio and maintain the pressure balance across the nuclear envelope, the relative nuclear size and mass have to increase (Figure 5L). Accordingly, nuclei occupy a higher percentage of the cell volume while the density ratio is maintained. These observations imply that the N/C volume ratio, in fact, is a consequence of cells maintaining a constant N/C density ratio with broad implications for cellular physiology (12).

**Figure 5.**
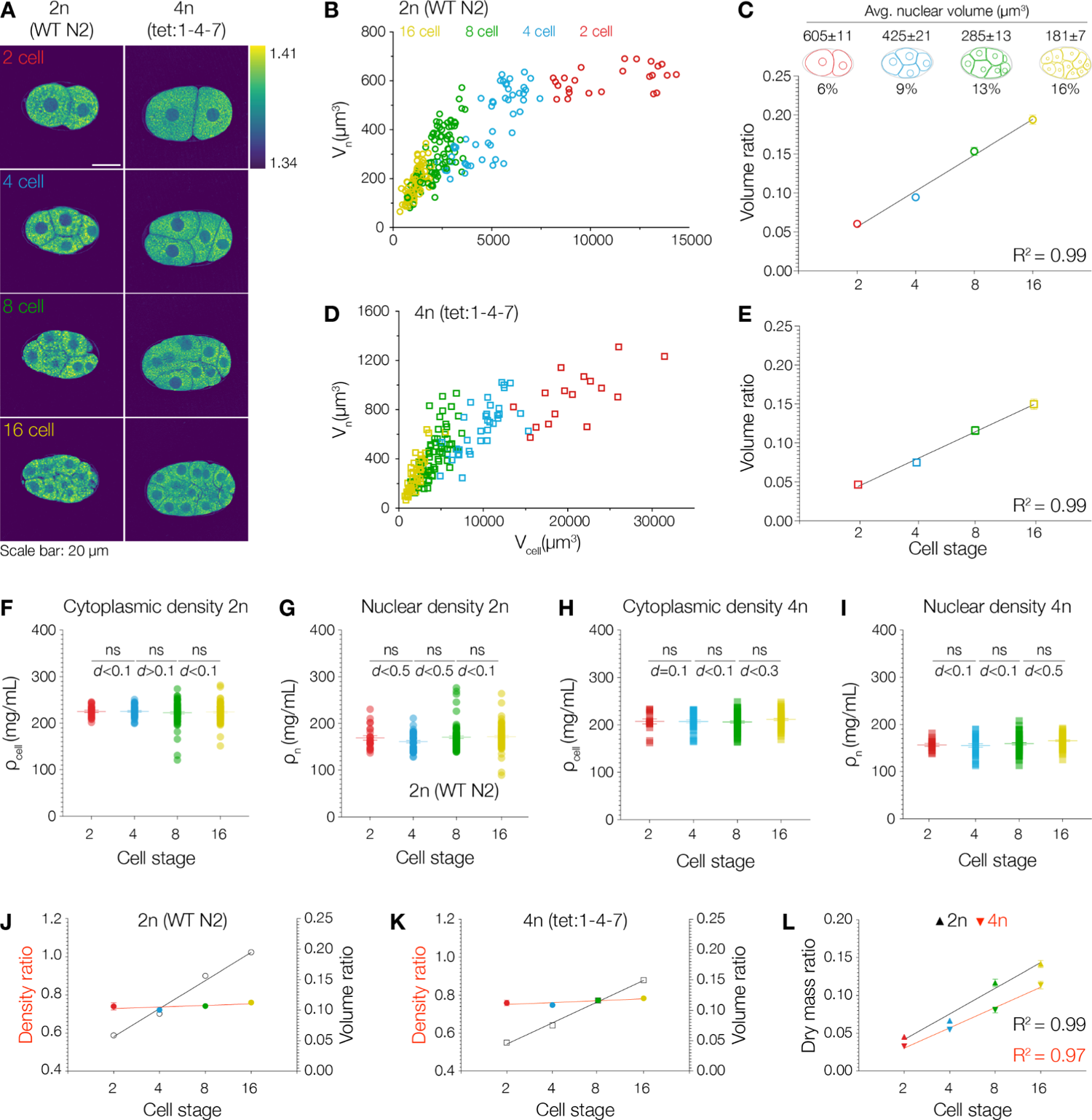
During early development, the nuclear-to-cytoplasmic density ratio is robustly maintained even when volume ratios change. (A) Representative ODT images of *C. elegans* embryos from different developmental stages. Left column shows wildtype N2 embryos (diploid) and tetraploid embryos. Nuclei in both wildtype and tetraploid embryos have a lower RI (and density) than the cytoplasm. Scale bar: 20 μm. Color bar on the right shows the RI distribution. (B) Nuclear volume scales with cell volume in wildtype embryos. (B-J) Red, blue, green, and yellow symbols represent values from 2 cell stage (n = 24), 4 cell stage (n = 48), 8 cell stage (n = 79) and 16 cell stage (n = 85) cells, respectively. (C) N/C volume ratios change during development in wildtype embryos. Average nuclear volumes (mean ± SEM) indicated above schematic for each stage and volume occupancy (% of nuclear volume/ cell volume, mean ± SEM) indicated below. Volume ratios at N/C_2Gell_ = 0.061 ± 0.002, N/C_4Gell_ = 0.095 ± 0.003, N/C8_Gell_ = 0.153 ± 0.005, and N/C_16oe_n = 0.194 ± 0.006. Bold black line indicates a linear fit of the data for the volume ratio. R^2^ value for fit indicated. (D) Nuclear volume scales with cell volume in tetraploid embryos. A similar power law relationship as in wildtype embryos is observed. 2 cell stage (n = 20), 4 cell stage (n = 44), 8 cell stage (n = 80), and 16 cell stage (n = 58) cells. (E) N/C volume ratios change during development in tetraploid embryos. Volume ratios at N/C2ce_l_i = 0.045 ± 0.002, N/C_4_c_eH_ = 0.074 ± 0.003, N/C_8_c_ell_ = 0.108 ± 0.005 and N/C_16_c_ell_= 0.150 ± 0.007. Bold black line indicates a linear fit of the data for the volume ratios. R^2^ value for fit indicated. (F) Cytoplasmic density is conserved through development in wildtype embryos. Densities (mean ± SEM) are ρ_cell_ = 225.3 ± 2.5 mg/mL, ρ_cell_ = 225.5 ± 1.9 mg/mL, ρ_cell_ = 222.5 ± 2.6 mg/mL, and ρ_cell_ = 224.1 ± 2.1 mg/mL. Each circle shows the value from a single cell. Mean ± SEM indicated. Mann-Whitney test where ns indicates p > 0.05. Cohen’s *d*2_G_e_l_&_4G_e_ll_< 0.1, Cohen’s *d*_4G_e_l_&_8G_e_ll_> 0.1, and Cohen’s *d*_8G_e_l_&1_6G_e_ll_< 0.1. (**G**) Nuclear density is conserved through development in wildtype embryos. Densities (mean ± SEM) are Pn_2cell = 169.1 ± 4.9 mg/mL, Pn_4cell = 161.1 ± 2.7 mg/mL, Pn_8cell = 170.7 ± 3.1 mg/mL, and Pn__1_6cell = 171.9 ± 2.6 mg/mL. Each circle shows the value from a single cell. Mann-Whitney test where ns indicates p > 0.05. Cohen’s *d*2_G_e_l_&_4G_e_ll_< 0.5, Cohen’s *d*_4G_e_l_&_8G_e_ll_> 0.5, and Cohen’s *d*_8c_e_ll_&1_6G_e_ll_< 0.1. (G) Cytoplasmic density is conserved through development in tetraploid embryos. Densities (mean ± SEM) are P2_G_e_ll_ = 204.6 ± 4.9 mg/mL, p_4_c_ell_ = 207.2 ± 3.4 mg/mL, ρ_cell_ = 206.2 ± 2.5 mg/mL, and ρ_cell_ = 211.4 ± 2.2 mg/mL. Each circle shows the value from a single cell. Mann-Whitney test where ns indicates p > 0.05. Cohen’s *d*2_G_e_l_&_4G_e_ll_= 0.1, Cohen’s *d*_4G_e_l_&_8G_e_ll_> 0.1, and Cohen’s *d*_8c_e_ll_&1_6G_e_ll_< 0.3. (H) Nuclear density is conserved through development in tetraploid embryos. Densities (mean ± SEM) are Pn_2cell = 156.4 ± 3.2 mg/mL, Pn__4_cell = 155.3 ± 3.3 mg/mL, Pn_8cell = 155.6 ± 2.4 mg/mL, and Pn__1_6cell = 165.2 ± 2.7 mg/mL. Each circle shows the value from a single cell. Mann-Whitney test where ns indicates p > 0.05. Cohen’s *d*2_G_e_l_&_4G_e_ll_< 0.1, Cohen’s *d*_4G_e_l_&_8G_e_ll_< 0.1, and Cohen’s *d*_8Gcell_&1_6Gcell_< 0.5. (I) Nuclear-to-cytoplasmic (N/C) density ratios are maintained even when N/C volume ratios change during development in diploid embryos. Graph shows the N/C density (circles) ratio at each stage. Density ratio at N/C_2cell_ = 0.74 ± 0.02, N/C_4cell_ = 0.72 ± 0.01, N/C_8_c_ell_ = 0.74 ± 0.01, and N/C_16cell_ = 0.76 ± 0.01. Red line indicates a linear fit of the density ratios. Black line with open circles indicates the data and a linear fit of volume ratios from panel (C). (J) Nuclear-to-cytoplasmic (N/C) density ratios are maintained even when N/C volume ratios change during development in tetraploid embryos. Graph shows the N/C density (circles) ratio at each stage. Density ratio at N/C_cell_ = 0.77 ± 0.01, N/C_4_c_ell_ = 0.76 ± 0.01, N/C_8cell_ = 0.77 ± 0.01, and N/C_16cell_ = 0.78 ± 0.01. Red line indicates a linear fit of the density ratios. Black line with open squares indicates the data and a linear fit of volume ratios from panel (E). (K) The nuclear dry mass ratio increases during development in both wildtype (triangles) and tetraploid (inverted triangles) embryos. For wildtype embryos dry mass ratio at N/C_2cell_ = 0.045 ± 0.002, N/C_4G_e_ll_ = 0.067 ± 0.003, N/C_8cell_ = 0.117 ± 0.005, and N/C_16cell_ = 0.142 ± 0.005. For tetraploid embryos dry mass ratio at N/C_cell_ = 0.033 ± 0.001, N/C_4cell_ = 0.055 ± 0.002, N/C_8cell_ = 0.081 ± 0.004, and N/C_16oe_|| = 0.114 ± 0.005. Bold black and red lines indicate a linear fit of the data for the wildtype and tetraploid, respectively. R^2^ values for each fit are indicated.

Taken together, our observations in *C. elegans* embryos show that N/C density ratios are maintained in the lineage of an embryonic system. They confirm our data from the *in vitro* reconstitutions in *Xenopus* and substantiate our mechanistic view on nuclear formation and growth. They further strengthen our pressure balance model which quantitatively links ploidy, nuclear dry mass, density, and volume.

## Discussion

While it is well appreciated that the physical and chemical properties of the cytoplasm and the nucleoplasm have far-reaching consequences for cellular function, subcellular density distributions remained largely unexplored. This study provides a mechanistic and quantitative understanding of how nuclei establish and maintain a lower density than the surrounding cytoplasm. Active import of nuclear proteins — assisted by entropic chromatin pressure — establishes a pressure difference across the nuclear envelope, which expands the nucleus to its final volume and resulting lower density. We show that a lower nuclear density with a constant N/C *density* ratio is robustly maintained even during early development when N/C *volume* ratios change. This implies that cells maintain a less crowded nucleus by adjusting their nuclear volume to cell volume. We find such a constant N/C density ratio of approximately 0.8 in 10 cell types ranging from yeast to human, which suggests that this ratio is a fundamental property ubiquitous to living systems.

In 1903 Richard Hertwig formulated the first quantitative hypothesis on organelle size (25). Based on his work on sea urchin embryos and algae, he proposed the “Kern-Plasma-Relation” as a constant characteristic for a given cell type (73, 74). Here, we identify a conserved ratio of nuclear-to-cytoplasmic density, which we propose to be the biophysical driver of the repeatedly observed “Kern-Plasma-Relation” (24, 42, 47–50, 58–61, 65, 70–80). Our experimental data together with the pressure balance model provide critical quantitative evidence for a mechanism which intrinsically links nuclear size to cell size by establishing a specific N/C density ratio. Furthermore, we provide several new and important biophysical quantities, e.g. nuclear and cytoplasmic density, dry mass, protein complex concentrations, and colloid osmotic pressures, which in the future can help constrain and advance many conceptual and theoretical models of nuclear size control and cellular density (10, 12, 47–50, 81).

In *Xenopus* egg extracts, our proposed model of pressure balance by localized nuclear proteins and chromatin sufficiently explains final nuclear size and density. It is, however, important to emphasize that using *Xenopus* egg extracts as a model system allows us to make simplifying assumptions. Cellular reality is likely more complex. Previous studies have characterized a multitude of cellular processes being involved in setting nuclear size. These include, among others, nucleocytoplasmic transport (42, 43, 58, 82, 83), gene expression and RNA processing (84), limiting cytoplasmic factors (42, 85–87), mechanical coupling between the cytoskeleton, the nuclear envelope, and chromatin (49, 88), and lipid homeostasis (89, 90). Thus, the proposed pressure balance will likely act in concert with these cellular processes and be constrained by other mechanical elements *in vivo* (91–94). Our model is qualitatively consistent with osmotic-based models of nuclear size scaling (13, 48–50) including the Pump-and-Leak model (48). How the parameters of our model will vary depending on the specific cellular system studied and, thus, contribute differentially to the quantitative pressure balance remains an important subject for future studies.

In this study, we explain how the unfolding and replicating chromatin contributes to the pressure balance both directly as a confined polymer-like chain and indirectly as a regulator of nuclear import. We thus propose that chromatin pressure has a modest but non-negligible direct effect on nuclear size. We model chromatin pressure as an entropic pressure. It has, however, been postulated that charged macromolecules including chromatin are surrounded by counterions to ensure electroneutrality (48, 49, 95). Collectively, these counterions could exert a substantial osmotic pressure. While some argue that under crowded conditions, osmotic pressures generated by counterions are negligible (46), our experiments do not allow us to discriminate between the purely entropic pressure of chromatin and the osmotic pressure exerted by chromatin counterions.

Finally, the question remains: why would it be important to maintain a robust density ratio between the nucleus and the cytoplasm? For one, it could simply be a physical consequence of the relative distributions of protein identities in the nucleus and the cytoplasm. We found absolute densities to be profoundly distinct (Figure S1B), this could imply that evolution shaped the cell-type specific proteome to optimize the trade-off between total protein concentration and colloid osmotic pressure (46, 96). Potentially though, a constant N/C density ratio evolved to serve a function. One conceptual idea is that a constant N/C density ratio is essential for a homeostatic coupling of transcription and translation. Indeed, it has recently been proposed that the maintenance of a homeostatic cell density is due to a scaling relation between amino acids, which are major osmolytes in the cell, and proteins, which contribute most of a cell’s dry mass (48). This is consistent with a dilute cytoplasm in senescent cells, which fail to scale nucleic acid and protein biosynthesis with cell volume (12), or with altered N/C volume ratios in cancer cells. Understanding how cell type-specific density and density distributions are set by balancing transcription, protein synthesis, and transport rates will remain an exciting avenue for future research.

## Materials and Methods

### Xenopus spec

The *Xenopus* frogs (adult females or males) used in this study are part of the *Xenopus* colony maintained at the animal husbandry of the Humboldt-Universität zu Berlin and were obtained from NASCO (Fort Atkinson, WI). *Xenopus* frogs were maintained in a recirculating tank system with regularly monitored temperature and water quality (pH, conductivity, and nitrate/nitrite levels) at 18-20°C (*X. laevis*) or 24-26 °C (*X. tropicalis*). All experimental protocols involving frogs were performed in accordance with national regulatory standards and ethical rules and reviewed and approved by the LaGeSo under Reg.-Nr. Reg 0113/20.

### C. elegans

Wildtype N2 strain worms (diploid) were picked and grown on NGM plates at 20°C. Single worms from strain SP346 (provided by the CGC (https://cgc.umn.edu/), which is funded by NIH Office of Research Infrastructure Programs, P40 OD010440) were picked and grown for several generations at 16°C on NGM/OP50 by selecting phenotypically long worms. Lines with consistent long phenotype and good fertility were selected. Oocyte chromosome numbers were counted to confirm tetraploidy of all lines. Line 1-4-7 was selected for all further experiments.

### S. cerevisiae

A haploid strain originally derived from the S288c strain (Genotype: *MATa SUC2 mal mel gal2 CUP1*) was used for imaging. S. cerevisiae were recovered from −80 °C glycerol stocks onto a 2% YPD plate. Colonies were inoculated into 5 mL SD media (0.17 % Yeast Nitrogen Base Without Amino Acids and Ammonium Sulfate, 0.5 % Ammonium Sulphate, 2 % Glucose) and incubated at 30 °C overnight with agitation. For imaging, cells were attached to imaging dishes (81156, μ-Dish 35 mm, ibidi) containing fresh SD media using Concanavalin A (C2010, Sigma; 1mg/mL).

### S. pombe

*S. pombe* strain AEP1 (Genotype: FY7385; h- leu1-32 ura4-D18 his3-D3) was grown in full medium (YES) at 30°C. For imaging, cells were transferred to imaging dishes (81156, μ-Dish 35 mm, ibidi) that were coated with Poly-L-Lysine (0.1% w/v, P6407, Sigma).

### C. reinhardtii

Motile *Chlamydomonas reinhardtii* wild-type strain CC125 were maintained in standard Tris-acetate phosphate (TAP) medium under continuous shaking at 110 rpm at 22°C. For imaging, cells were transferred to imaging dishes (81156, μ-Dish 35 mm, ibidi) that were coated with Poly-L-Lysine (0.1% w/v, P6407, Sigma).

### D. melanogaster

*Drosophila* S2R+ cells were maintained in Schneider’s medium (11720-034, Gibco) at 23°C and 33% relative humidity. Cells were detached from T-75 tissue culture flasks by thoroughly pipetting the medium within the flask. For imaging, cells were allowed to attach to imaging dishes (81156, μ-Dish 35 mm, ibidi) that were pre-coated with Poly-L-Lysine (0.1% w/v, P6407, Sigma).

### D. rerio

Fertilized embryos were obtained from AB strain zebrafish. Embryos were allowed to grow for 4 hours post fertilization in E3 medium. Embryos at the dome stage 4-5 were collected and dissociated by gently pipetting up and down in deyolking buffer (55 mM NaCl, 1.8 mM KCl, 1.25 mM NaHCO_3_ in HBSS, Life Technologies). For imaging, dissociated embryonic cells were transferred to imaging dishes (81156, μ-Dish 35 mm, ibidi).

### M. musculus

R1/E mouse embryonic stem cells were cultured as in (97). Briefly, cells were maintained in DMEM media (41966-029; Gibco) supplemented with 16% FBS (Gibco), antibiotic-antimycotic (Invitrogen), nonessential amino acids (Gibco), P-mercaptoethanol (Gibco), and recombinant mouse leukemia inhibitory factor (ESGRO). Cells were passaged every 48 h and seeded at a density of 35,000 cells/cm^2^ onto gelatin-coated dishes. For imaging cells, were transferred to imaging dishes (81156, μ-Dish 35 mm, ibidi) that had been coated with Laminin-511 (2.5 μg/mL, LN511, BioLamina)

### sapiens

HeLa S3 cell lines were cultured in DMEM (41966-029; Gibco) supplemented with 10% FBS and antibiotic-antimycotic (Invitrogen) in an incubator maintained at 37°C with 5% CO_2_ and passaged routinely. For imaging, cells were transferred onto imaging dishes (81156, μ-Dish 35 mm, ibidi).

### Imaging and image analysis

#### Correlative optical diffraction tomography and confocal fluorescence microscopy

Refractive index (RI) tomograms were obtained using a custom-built optical diffraction tomography (ODT) setup based on Mach-Zehnder interferometry. The setup was built onto a commercial inverted microscope stand (IX81, Olympus Life Science), which increased usability and allowed us to use separate light paths for ODT, epifluorescence, and confocal fluorescence microscopy (Figure S1). The detailed description of a similar ODT setup can be found in (16, 28). Briefly, a coherent laser beam (λ = 532 nm) was split into two using a single-mode fiber optic coupler. One beam was used as a reference beam and the other was used to illuminate the sample using a tube lens (f = 175 mm) and a 60x water-dipping objective lens (LUMPLFLN60XW, NA 1.0, Olympus Life Science). To reconstruct 3D RI tomograms, the sample was illuminated from 150 different incident angles by a dual-axis galvanometer mirror (GVS012/M, Thorlabs Inc.) covering the azimuthal angular range from −48° to 48°. The scattered light from the sample was collected by a high numerical aperture 100x objective lens (oil immersion, UPlanFl, NA 1.3, Olympus Life Science) and interfered with the reference beam at the image plane, which generated spatially modulated holograms. The holograms were recorded by a CCD camera (FL3-U3-13Y3M-C, FLIR Systems, Inc.). The detailed principle for tomogram reconstruction can be found in (98, 99). Briefly, from the spatially modulated holograms, the complex optical fields of light diffracted by the sample were retrieved by applying a field retrieval method based on Fourier transformation (100). The 3D RI tomogram of the sample was reconstructed by mapping 2D Fourier spectra of the retrieved complex optical fields onto the surface of Ewald spheres corresponding to the spatial frequency of the incident angles based on Fourier diffraction theorem (98–102). The Gerchberg-Papoulis (GP) constraint is applied to fill the missing cone artifact.

Confocal fluorescence microscopy was performed using a Rescan Confocal Microscope (103, RCM1 from confocal.nl) which was installed on the same microscope frame. Fluorescence images or image stacks (with a step size of 1 μm) were acquired using the same high numerical aperture objective lens (UPlanFl, NA 1.3, Olympus Life Science) and recorded on to a CMOS camera (CM3-U3-50S5M-CS, FLIR Systems, Inc. or FL-AB-20-BW Tucsen Photonics Co., Ltd.). The RCM1 and microscope frame was controlled using a custom Micromanager script (provided by confocal.nl).

#### Calculating mass densities from refractive index values

3D refractive index (RI) tomograms were reconstructed using a custom Matlab script (16, 28). The 3D tomogram was computed based on Fourier diffraction theorem (102) from multiple 2D quantitative phase images that were acquired by illuminating the sample at various oblique angles. Upon tomogram reconstruction each voxel within the 3D RI tomogram has an assigned RI value. The absolute RI values for the surrounding medium (ni, where i = extract or medium) were independently measured using an Abbe refractometer (Arcarda ABBE-2WAJ). Since the RI value in most biological samples is linearly proportional to the mass density of material with the proportionality coefficient, or RI increment, α (104), mass densities within each voxel were calculated using the relationship n_sample_ = ni + ap_sample_, where α is the refractive index increment for proteins/nucleic acids (0.19 mg/mL; 31) and p_sample_ is the mass density within the sample. For control extract samples, the mass density of ni (refractive index of the surrounding cytoplasm) was measured to be 100 mg/mL (based on protein concentration measurements using the Bradford assay). During perturbation experiments, the mass density value of ni was determined based on the measured protein concentration of the sample (such as for HS extracts, during osmotic shock or upon adding BSA). For measurements in buffer and cell culture medium, the RI of the medium (ni) was measured using an ABBE refractometer and used as a background reference.

#### Image analysis and quantification

Nuclear/cytoplasmic region of interest (ROI) determination and nuclear volume segmentation was performed using FIJI (105). Within cells from different species, nuclear and cytoplasmic mass densities were measured using square ROI (0.5 μm length for *S. cerivasiae* and *S. pombe*, 1 μm length for *C. reinhardtii*, 5 μm length for *C. elegans* and *D. rerio*, 2 μm length for *D. melanogaster* and *X. laevis*, and 2.5 μm length for *M. musculus* and *H. sapiens*). Arbitrary and unbiased selections were made for each nuclear and cytoplasmic ROI. For nuclei containing nucleoli, nucleolar regions were excluded from the analysis of nucleoplasmic mass density. For *Xenopus* egg extracts, nuclei were segmented from the 3D tomograms either manually or using the 3D volume manager plugin (part of the Bio Image Analysis Toolbox from the SCF, MPI-CBG; https://sites.imagej.net/SCF-MPI-CBG/). To calculate nuclear volumes, the voxels within the segmented regions were integrated. To calculate nuclear dry mass, the mass densities within each segmented volume were integrated. To calculate the average mass density within each segmented nucleus, the total dry mass was divided by the total volume. To quantify fluorescence intensities (Hoechst-33342/NLS-GFP) nuclei within images were segmented either manually or using Otsu thresholding. To calculate fluorescence intensities, the gray values within the segmented regions were integrated. Next, the background intensities were calculated for regions outside nuclei with dimensions similar to the segmented nuclear regions. Finally, the total fluorescence intensity within the sample was divided by the total background intensity and these values were reported in the figures.

#### C. e*legans* embryo ODT imaging and analysis

Custom-built ODT setups were used to image wildtype and tetraploid embryos. Embryos were isolated from healthy adults and placed onto coverslips (631-1573P, VWR International). A drop of M9 buffer (∼10 μL) containing 20 μm polystyrene beads was placed on the coverslip along with the embryos. A second smaller coverslip (631-0124, VWR International) was gently placed on top of the embryo to create a confined volume for imaging which reduced scattering. The beads acted as spacers and prevented squashing of the developing embryos.

For *C. elegans* embryos in addition to the regular field retrieval and tomogram reconstruction code, an axial drift during the time lapse imaging was corrected by identifying the center of mass of the reconstructed RI tomograms. Further, since the embryos were placed between 20 μm beads, axial constraints were implemented within the code to reduce scattering and improve the quality of the reconstructed tomograms. Embryos and nuclei were segmented using the 3D volume manager plugin and the segmented masks were used for calculating volumes. The average mass density within the segmented volumes were reported and total dry mass was calculated by integrating the mass density information over the 3D volume. The N/C volume ratio was calculated by dividing the nuclear volume by the cytoplasmic volume (cell volume - nuclear volume). The N/C density ratio was calculated by dividing the average mass density of the nucleus by the average mass density of the cytoplasm. The dry mass ratio was calculated by dividing the total nuclear dry mass by the total cytoplasmic dry mass.

#### *C. elegans* brightfield and fluorescence imaging

For bright field imaging, worms were imaged with an Apo Z 1.5x objective and BF+ light on a Zeiss Axio Zoom V16 dissection scope equipped with an Axiocam 705 mono camera and Zen 3.2 software. Embryos were imaged with bright field illumination on a Leica DM6000B microscope with a 63x.1.30 HC PL Apo glycerol objective and an Andor iXon Life camera using Visiview (Visitron) software.

To confirm tetraploidy of the 1-4-7 strain, 1% polylysine solution was spotted onto microscope slides and dried. 4 μL M9 buffer were added and 20 adult worms were picked into the drop under a dissection scope. Buffer was carefully removed with a strip of Kimwipes and worms stuck to the polylysine surface. 4 μL of 95% ethanol were added and desiccated. 4 μL 95% Ethanol drops were added again and desiccated and this cycle was repeated four times. 10 μL of Vectashield with 1 μg/mL Hoechst-33342 was pipetted onto the desiccated worms and they were allowed to rehydrate. A 22 mm coverslip was gently placed onto the drop and nail polish was used to seal the coverslip. Fluorescence images of oocyte chromosomes in utero were taken in the DAPI channel on a DeltaVision RT imaging system (Applied Precision, LLC; IX70 Olympus) equipped with a charge-coupled device camera (CoolSNAP HQ; Roper Scientific) in 33 x 0.7 μm z-sections using an Olympus 63×1.43 NA UplanSApo objective. Image stacks were deconvolved using Softworx (Applied Precision, LLC) and maximum-intensity projected for oocyte chromosome planes with FIJI (105). For counting chromosomes, individual z-planes were assessed in cases of overlapping chromosomes in the maximum-intensity projections.

#### FRET imaging

To measure RanGTP levels in reconstituted nuclei, a YRC probe (final concentration 2 μM) was added to nuclear assembly reactions. Nuclei were sandwiched between coverslips and images were acquired of the CFP and FRET channels. Imaging was performed on a Nikon Widefield Ti2 fluorescence microscope equipped with an sCMOS camera (PCO. edge) and a 40x water immersion objective lens (CFI Apo Lambda S 40XC WI). The system was controlled using the Nikon Elements AR software. Samples were illuminated using LEDs (Lumencor, SpectraX). A 200 ms exposure time was used for acquiring the FRET and CFP images. Image analysis was performed using FIJI (105). Briefly the FRET and CFP images were divided to obtain an intensity ratio image containing the I_FR_ET/I_CFP_ values. To quantify intensity profiles, line scans (line width: 30 pixels) were performed over a 30 μm length along the major axis of nuclei.

#### Nuclear assembly in *Xenopus* egg extracts

Metaphase-arrested egg extract was prepared from laid *X. laevis* eggs as previously described (106, 107). Briefly, *X. laevis* frogs were primed with 100 U of pregnant mare serum gonadotrophin (PMSG) 3-7 days before the experiment and were boosted with 1000 U human chorionic gonadotrophin (HCG) to induce egg laying. Eggs arrested in the metaphase stage of meiosis II were collected, dejellied using L-Cysteine and fractionated via centrifugation. The cytoplasmic layer was then isolated and supplemented with Cytochalasin and Complete EDTA-free protease inhibitor. Nuclei were assembled in extracts similar to previously published protocols (33). Extracts were cycled to interphase using CaCl2 (final concentration: 0.6 mM). Cycloheximide (final concentration: 0.1 mg/mL) and energy mix were added to arrest extracts in interphase and support nuclear assembly, respectively. To visualize DNA, membranes and assay nuclear import Hoechst-33342 (final concentration: 0.05 mg/mL), Dil (final concentration: 0.1 mg/mL) and NLS-GFP (final concentration: 0.2 mg/mL) were added. 100 μL extract reactions were prepared for each experimental test. Finally, demembranated sperm nuclei (final concentration: 1000/μL) were added to each tube and incubated in a water bath maintained between 16-20°C. Extracts were mixed every 15 minutes with the help of a precut pipette tip.

To compare the effects of altering the chromatin content on nuclear size, nuclei were assembled from either *X. laevis* or *X. tropicalis* sperm using the same batch of *X. laevis* interphasic extract. *X. tropicalis* sperm nuclei were obtained from adult *X. tropicalis* male frogs as described previously (108, 109).

For imaging, nuclei were sandwiched between two coverslips (631-1573P, VWR International) at different time points. Another coverslip was immediately placed on this coverslip to create a squashed egg extract sample, which provided a thin layer that could be easily imaged with our imaging setup.

#### Protein concentration measurements

Bradford reagent (B 6916, Sigma) was used to determine the concentration of proteins in egg extract and other solutions in this study. As per the manufacturer’s recommendation a protein standard curve was first measured using Bovine Serum Albumin (P 0384, Sigma) standards. The standards were dissolved in CSF-XB buffer. To measure protein concentration via the Bradford method, extract was first diluted (100 or 200-fold) in CSF-XB buffer. Next, 50 μL of the protein solution were added to an Eppendorf tube along with 1.5 mL Bradford reagent. Samples were incubated at room temperature for 15 minutes and subsequently transferred to polystyrene cuvettes (Y195.1, Roth) for absorption measurements. Absorption at 595 nm was measured using an Eppendorf BioSpectrometer Basic. Protein concentrations were determined by matching the absorbance values with the concentration information using the standard curve.

#### Protein purification

The GST-NLS-GFP plasmid (pMD49) was a gift from Dr. Thomas Quail (EMBL Heidelberg) and the GST-NLS-GFP protein was purified as described previously (42). The YFP-RBD-CFP (YRC) chimera protein plasmid (pKW966) was a gift from Prof. Karsten Weis (ETH Zürich), the protein was purified as described previously (55). Recombinant Nucleoplasmin (Npm2) was purified as described (39).

#### Nucleoplasmin immunodepletion and add-back

To investigate whether Nucleoplasmin2 (Npm2) was necessary for reducing mass density, sperm nuclei in buffer (containing energy mix) were supplemented with recombinant Npm2 (final concentration: 9 μM). The sample was incubated between 16-20°C and imaged at 10 and 30 minutes after the addition of Npm2. To immunodeplete Npm2 from *Xenopus* egg extract purified polyclonal Npm2 antibody (rabbit α-Npm2 IgG, 39) were first cross linked to Protein A beads (Dynabeads™ Protein A, 10001D, ThermoFisher) using Dimethyl pimelidate dihydrochloride (D8388, Sigma). 80 μg of Npm2 antibody were cross-linked to 330 μl of Protein A beads (10001D, ThermoFisher Scientific). Interphase *Xenopus* egg extract containing cycloheximide was prepared as described above and immunodepletions were performed in a cold room. The extract was subjected to two rounds of immunodepletion and protein reduction levels were assayed using Western blots. Along with immunodepleted samples, a control depletion was performed using the same volume of empty Protein A beads. To test if Npm2 was sufficient for reducing mass density, purified Npm2 was added back to immunodepleted extract to physiological concentrations (4.2 μM, 85).

#### Protein gels and Western blots

Protein samples from each condition were supplemented with SDS buffer and boiled at 95°C for 10 minutes. Samples were centrifuged at 1600 g for 5 minutes. The supernatant was loaded into wells of a protein gel (NP0301BOX, Invitrogen) with SDS running buffer (NP0001, Invitrogen). Protein bands were separated by running the gel and Instant Blue Coomassie stain (ab119211, abcam) was used to visualise bands. Protein bands were transferred to a PVDF membrane (8858, ThermoFisher) using a transfer buffer (NP0006, Invitrogen). Membranes were blocked using a blocking buffer (5% dry milk in TBST) for 60 minutes at RT with gentle rocking. Next, the blots were incubated with the primary Npm2 antibody (1:10,000 dilution in blocking buffer) overnight at 4°C with gentle rocking. A monoclonal αTubulin-antibody (1:15,000 dilution in blocking buffer; T9026, Sigma) was used as a loading control. Blots were washed thrice with TBST with 5 minute incubations each. Next the secondary antibody (1:5000 in blocking buffer; anti rabbit-HRP for Npm2 and anti mouse-HRP for αTubulin) was added and incubated for 1 hour at room temperature with gentle rocking. The blots were finally washed thrice with TBST with 5 minute incubations before adding the ECL substrate solution (1705061, Biorad). Chemiluminescence images were acquired on a Chemidoc™ imaging system (Biorad). Band intensities were quantified using FIJI (105). Total intensities from the monomeric and pentameric bands were added and normalised to the intensity of the loading control bands.

#### Biochemical perturbations

For biochemical perturbations, inhibitors and drugs were added to the interphasic extracts before the addition of sperm nuclei and the initiation of nuclear assembly. Stock solutions were prepared to ensure that addition of inhibitors did not dilute the extract by more than 10% of the original extract volume. To perturb import, a small molecule inhibitor cocktail of Ivermectin (I8898, Sigma; final concentration: 100 μM) and Pitstop-2 (SML1169, Sigma; final concentration: 30 μM) was used. To prevent chromatin decondensation an inhibitor cocktail of the DNA intercalator Actinomycin D (A1410, Sigma; final concentration: 10 μg/mL), topoisomerase II inhibitor ICRF-193 (I4659, Sigma; final concentration: 150 μM) and the calcium chelator BAPTA (196418, Sigma; final concentration: 5 mM) was supplemented to interphase extracts. To inhibit DNA replication, Aphidicolin (A0781, Sigma; final concentration: 200 μM) was added to the extract before nuclear assembly. To reduce or increase protein concentrations, extracts containing pre-assembled nuclei were either diluted using CSF-XB buffer.

#### Osmolality measurements

A freezing point osmometer (Osmomat 3000 Basic, Gonotec) was used for measuring the osmolality of different extract and protein solutions. 50 μL of sample was used for each assay and at least 2 independent measurements were made for each solution.

#### Quantification and statistical analysis

Details about the quantifications are provided in each figure legend, including the total number of observations (n), the mean, and SEM values. Further statistical tests used for measuring significance, effect size, and their interpretation is provided. For statistical analysis and plotting, we utilized GraphPad Prism version 9.0 for Mac OS X, GraphPad Software, La Jolla California USA, www.graphpad.com. The alpha value was set at 0.05 and the p value was calculated to test how different the groups were from each other. P values greater than 0.05 are represented by ‘‘ns’’. A single * indicates a p value < 0.05, ** indicates p values < 0.01, *** indicates p values < 0.001, and **** indicates p values < 0.0001. In addition to the significance, we also indicate the effect size by calculating Cohen’s d. An effect size between 0.20 - 0.50 was considered small, while effect sizes between 0.51 - 0.80 were considered medium and d > 0.81 was considered large. For the linear fits, bold lines indicate the best fit and thin/dotted lines indicate the 95% confidence interval. R^2^ values indicated for each fit. When necessary, graph visuals such as line thickness, fonts, and colours were optimised using Adobe Illustrator.

## Author contributions

This work represents a truly collaborative effort. Each author has contributed significantly to the findings and regular group discussions guided the development of the ideas presented here. In more detail, A.B. performed all experiments and analyzed the data. O.M. and V.Z. conceptualized and wrote the theory part. K.K. designed the ODT setup and wrote the script for controlling the setup. A.B. and K.K. built the correlative fluorescence-ODT setup. K.K. developed the script for field retrieval, tomogram reconstruction and axial drift correction. A.B., K.K. and C.H. performed the *C. elegans* experiments. B.L. and D.S. purified the recombinant Npm2 and Npm2 antibodies. O.M., K.K., V.Z., D.S. and J.G. contributed to experimental design, data analysis, and interpretation. J.G. V.Z. and S.R. conceived the project. S.R. wrote the manuscript with input from all authors. The authors declare no competing interests.

## Acknowledgements

The authors would like to thank all current and former members of the Guck, Reber, and Zaburdaev labs. We thank Dr. Roland Knorr (HU Berlin) and Dr. Alexander May (Tokyo Institute of Technology) for the S*. cerevisiae* cells, Auguste Palm (FU Berlin) and Prof. Dr. Ann Ehrenhofer Murray (HU Berlin) for the *S. pombe* cells, Dr. Olga Baidukova and Prof. Dr. Peter Hegemann (HU Berlin) for the *C. reinhardtii* cells, Dr. Alf Herzig (MPIIB-Berlin) for the *D. melanogaster* cells, Anika Neuschulz and Dr. Jan Philipp Junker (MDC Berlin) for the *D. rerio* cells. We would like to thank Dr. Thomas Quail (EMBL Heidelberg) and Prof. Dr. Karsten Weis (ETH Zurich) for the GST-NLS-GFP and YRC plasmids, respectively. We also thank Tobias Kletter (MPI-IB) for help with figure design, Prof. Dr. Simon Alberti and Dr. Anna Taubenberger (TU-Dresden) for permission to use their correlative fluorescence-ODT setup, and Dr. Jan Schromanzer and his team at the Advanced Medical BIOimaging Core Facility, Charité Berlin. We thank the labs of Matthew Kraushar (MPI-MG) and Marcus Taylor (MPI-IB) for ongoing discussions and critical reading of the manuscript, and Dirk Görlich (MPI-Nat), Mathieu Piel and Romain Rollin (Institut Curie) for helpful comments on the manuscript.

## Supplemental information

### Supplemental figures S1-S6

**Figure S1.**
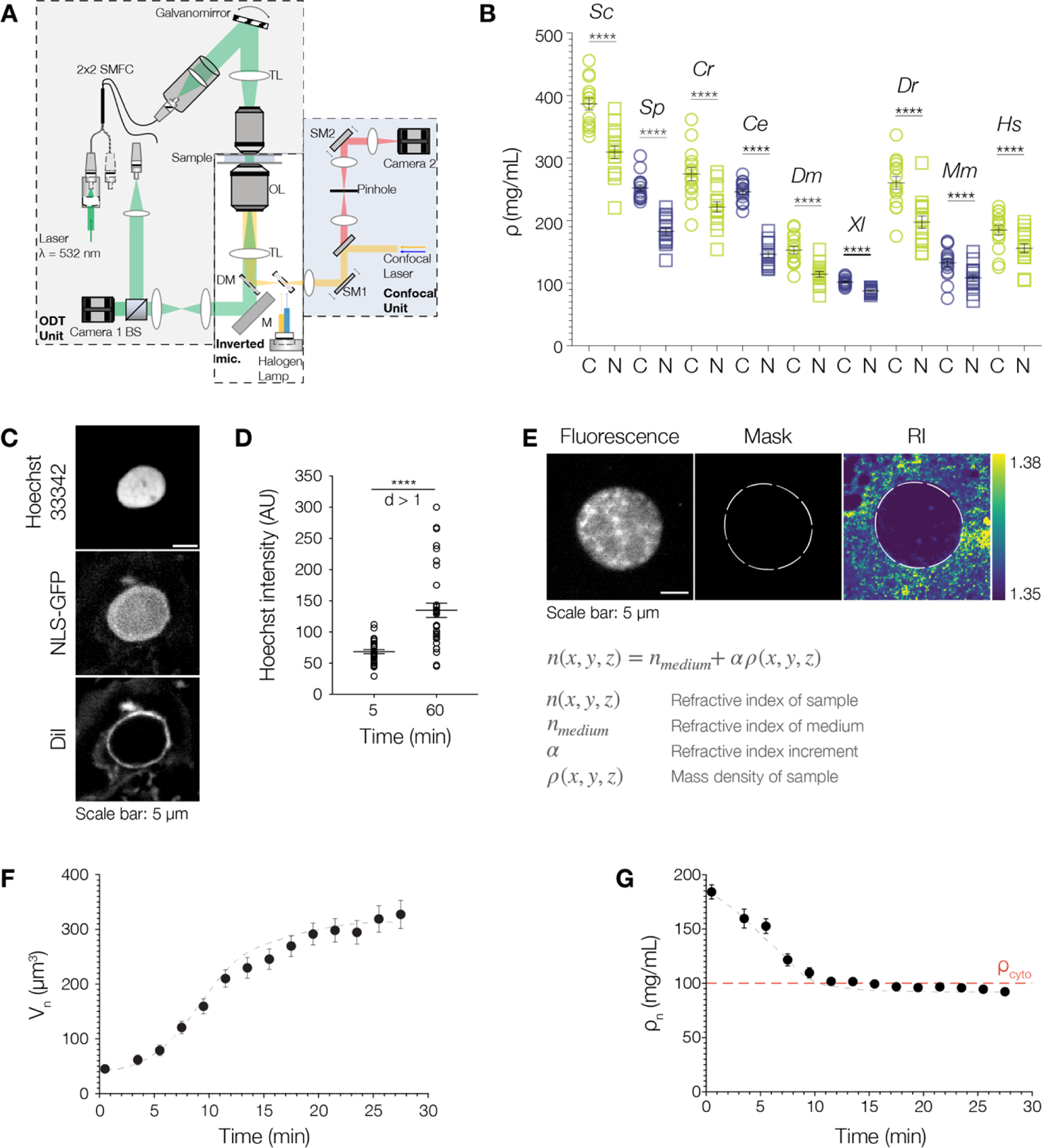
A combined confocal fluorescence microscopy and optical diffraction tomography setup provides volume and density information during nuclear assembly in *Xenopus* egg extracts. (A) Schematic of the optical setup used for experiments. SMFC, single mode fibre coupler; TL, tube lens; CL, condenser lens; OL, objective lens; M, mirror; DM, dichroic mirror; BS, beam splitter; SM1, scanning mirror 1; SM2, scanning mirror 2. See Materials & Methods for more details. (B) Quantification of the cytoplasmic (C, green circles) and nuclear (N, blue squares) density (ρ) across cells from different organisms. *Sc*: *S. cerevisiae*, *Sp*: *S. pombe*, *Cr*: *C. reinhardtii*, *Ce*: *C. elegans*, *Dm*: *D*. *melanogaster, Xl*: *X. laevis, Dr*: *D. rerio, Mm*: *M. musculus* and *Hs*: *H. sapiens*. n = 15 measurements from randomly selected ROIs in the cytoplasm and nucleus, respectively. Nuclear densities were measured in nucleoplasmic regions excluding nucleoli. Bars indicate the mean ± SEM in all graphs. Mann-Whitney test where **** indicates p < 0.0001. (C) Nuclei assembled in *Xenopus* egg extract are import competent. Representative fluorescence image where DNA was stained with Hoechs-33342, NLS-GFP was used to show active import and membranes were stained with Dil. (D) Nuclei assembled in *X. laevis* egg extracts are replication competent as indicated by the quantification of total Hoechst-33342 intensity within assembling nuclei at 5 minutes (n = 31) and 60 minutes (n = 33). The average intensity values double from 68.6 ± 3.2 AU to 134.8 ±11.5 AU. Bar indicates the mean ± SEM. Mann-Whitney test where **** indicates p < 0.0001. Cohen’s *d*_5_&_60min_>1. (E) Representative fluorescence image of a *Xenopus* nucleus (stained with Hoechst-33342) at 60 min and corresponding mask obtained by segmentation. Masks were obtained from the fluorescence images and transferred to the ODT images to quantify the mass density of regions of interest. Scale bar = 5 μm. Color bar on the right shows the RI range in the ODT image. Equation used to obtain the mass density (ρ) of the sample from the 3D RI using the refractive index increment *α*. See Material & Methods for more details on converting refractive index to mass density. (F) Quantification of nuclear volume at 2 min time intervals as nuclei assemble. (G) Quantification of nuclear mass density at 2 min time intervals as nuclei assemble. Red line shows the mass density of the surrounding cytoplasm (ρ_cyto_). For (F) and (G), n = 38 nuclei from 3 independent experiments, circles represent mean and bars represent the SEM.

**Figure S2.**
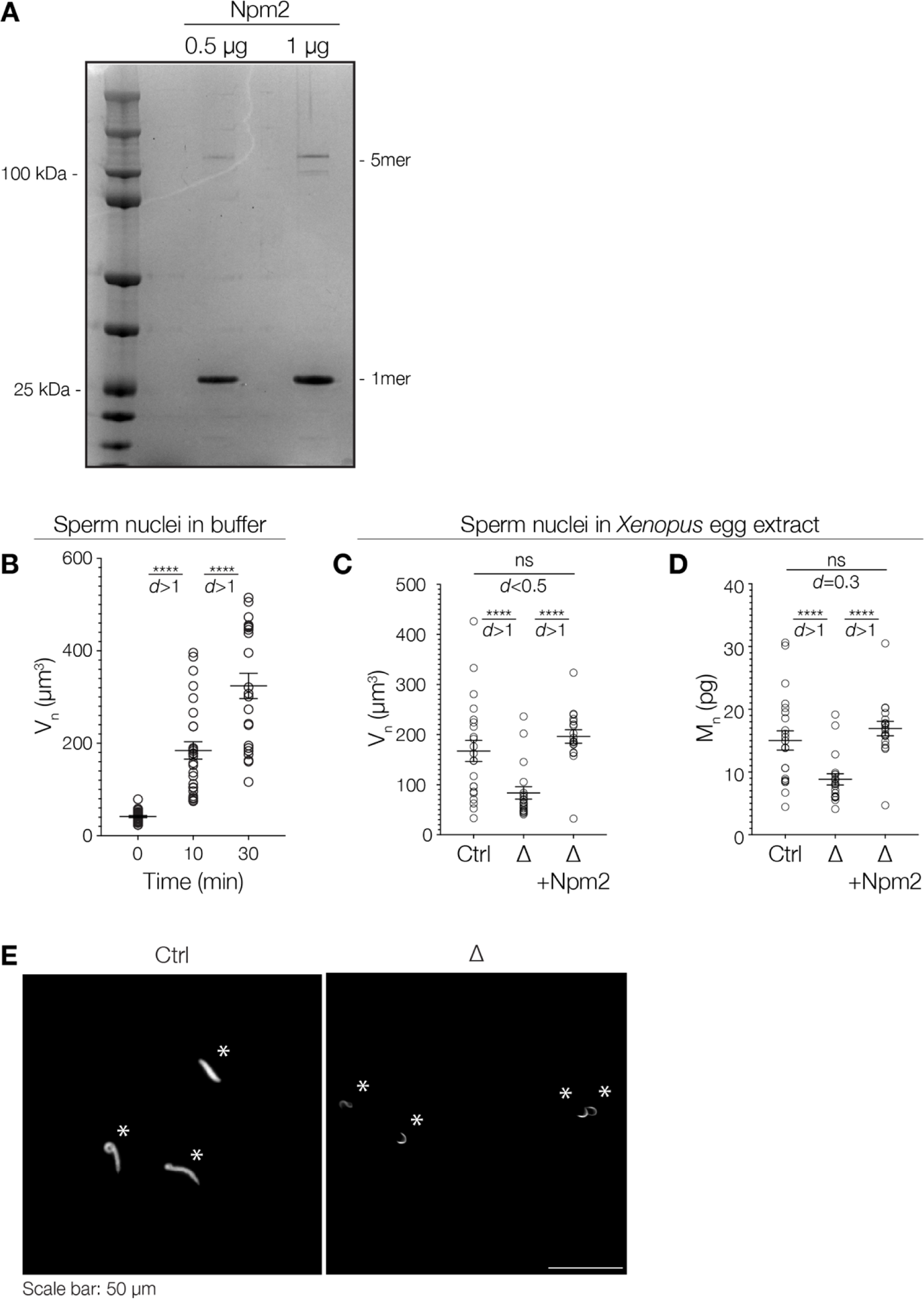
Nucleoplasmin is both necessary and surrounding solvent. (A) Coomassie stain of an SDS gel showing purified the pentameric and monomeric form of the protein. lane shows the molecular weight marker. sufficient to reduce nuclear density to that of the Npm2 protein. Two distinct bands can be seen for 0.5 pg and 1 pg of protein were loaded. Left most (B) Chromatin volume in buffer at 0, 10, and 30 minutes after the addition of Npm2 (n = 30, 27, and 23, respectively from 3 independent experiments). Volume (mean ± SEM) V_n_0_ = 41 ± 2 μm^3^, at V_n_10_ = 184 ± 19 μm^3^ and at V_n_30_ = 324 ± 27 μm^3^. Mann-Whitney test where **** indicates p < 0.0001. Cohen’s *d*_0_&_10min_ > 1 and Cohen’s *d* _10_&_30min_> 1. (C) Chromatin volume in control-depleted, ANpm2 and A+Npm2 extracts. Sperm nuclei in ΔNpm2 extracts have a lower volume (84 ± 12 μm^3^) than decondensing sperm nuclei in control-depleted (167 ± 21 μm^3^) and Δ+Npm2 (196 ± 13 μm^3^) extracts (n = 19, 22, and, 18 samples from 2 independent experiments) after 10 minutes. There is no significant difference between the volume of sperm nuclei in control-depleted and Δ+Npm2 extracts. Bars show mean ± SEM. Mann-Whitney test where ns indicates p > 0.05 and **** indicates p < 0.0001. Cohen’s *d*ct_rl&ANpm2_ > 1, Cohen’s *d*_ANpm2_&A_+Npm_2 > 1 and Cohen’s *d*MOCK&A+Npm2 < 0.5. (D) Quantification of dry mass in control-depleted, ANpm2 and A+Npm2 extracts. Sperm nuclei in ΔNpm2 extracts have a lower dry mass (8.8 ± 0.9 pg) than decondensing sperm nuclei in control-depleted (15.1 ±1.5 pg) and Δ+Npm2 (16.9 ± 1.1 pg) extracts (n = 19, 22, and 18 samples from 2 independent experiments) after 10 minutes. There is no significant difference between the dry mass of sperm nuclei in control-depleted and Δ+Npm2 extracts. Bars show mean ± SEM. Mann-Whitney test was used to test statistical significance where ns indicates p > 0.05 and **** indicates p < 0.0001. ^Cohen s^ *d*Ctrl&ANpm2 > ^1^, ^Cohen s^ *d*ANpm2&A+Npm2 > ^1^ and ^Cohen s^ *d*MOCK&A+Npm2 = 0.3. (E) Low magnification fluorescence images showing sperm nuclei (asterisks) stained with Hoechst-33342 in control-depleted and ΔNpm2 extracts after 10 minutes of incubation. Scale bar: 50 pm.

**Figure S3.**
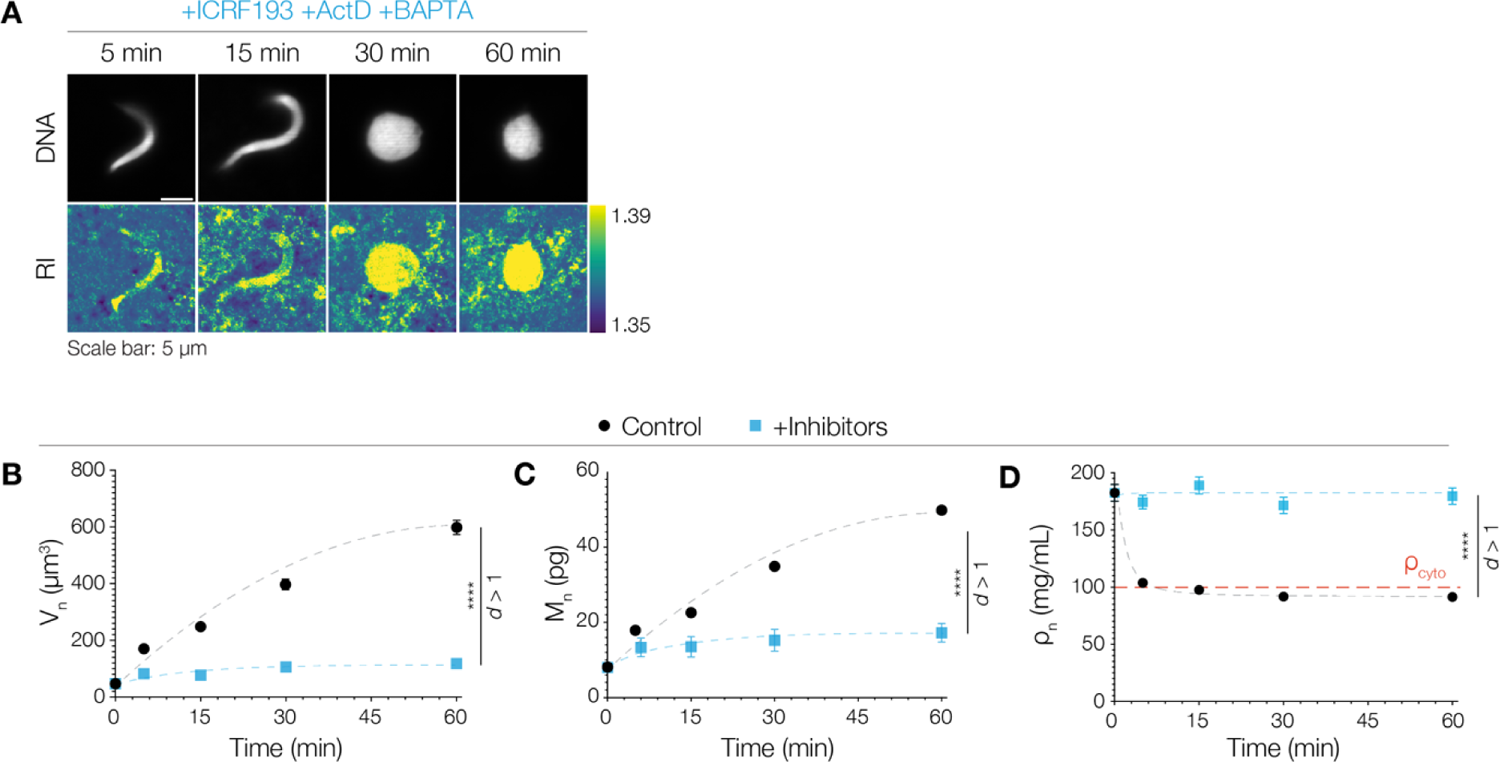
Inhibition of chromatin decondensation and nuclear import leaves nuclei with a density comparable to that of fully condensed sperm chromatin. (A) Nuclei were assembled in *Xenopus* egg extracts in the presence of ICRF193 (a topoisomerase inhibitor), Actinomycin-D (ActD, a DNA intercalator) and BAPTA (a calcium chelator). This inhibitor cocktail inhibits chromatin decondensation and allows for poreless nuclear envelope closure. Top panel shows representative fluorescence images of Hoechst-33342 stained nuclei (DNA) from different time points. Bottom panel shows the corresponding RI (central slice from the ODT) image. While assembling nuclei round up, they have a RI value comparable to that of fully condensed sperm chromatin. (B) Volume of poreless nuclei. V_n_ (blue squares, mean ± SEM, n = 20 from 2 independent experiments) at indicated time points: V_n__5 = 80 ± 10 pm^3^, V_n_15_ = 75 ± 10 pm^3^, V_n_30_ = 95 ± 19 pm^3^ and V_n_60_ = 105 ± 12 μm^3^. Black circles show mean at different time points for control nuclei. At 60 minutes there is a significant difference between the volumes of nuclei. Mann-Whitney test where **** indicates p < 0.0001. Cohen’s *d*Contro&+InhiMors> 1. In panels (B-D). (C) Dry mass of poreless nuclei. Blue squares show mean at different time points for +Inhibitors nuclei. M_n_ (blue squares, mean ± SEM, n = 20 from 2 independent experiments) at M_n_5_ = 13.5 ± 1.9 pg, M_n_15_ = 13.5 ± 2.1 pg, M_n_30_ = 15.2 ± 2.2 pg and M_n_60_ = 17.2 ± 2.1 pg. Black circles show the values at different time points for control nuclei. Mann-Whitney test where **** indicates p < 0.0001. Cohen’s *d*_Control_&_+Inhibitors_ > 1. (D) Density of poreless nuclei. ρ_n_ (blue squares, mean ± SEM, n = 20 from 2 independent experiments) at indicated timepoints: p_n_5min_ = 174.4 ± 6.0 mg/mL, p_n_i5min_ = 189.0 ± 7.4 mg/mL, p_n_30min_ = 171.5 ± 7.1 mg/mL and p_n_60min_ = 179.6 ± 7.2 mg/mL. For poreless nuclei, p_n_ does not change significantly during nuclear assembly. Black circles show mean at different time points for control nuclei. Red line: ρ of the cytoplasm. Mann-Whitney test where **** indicates p < 0.0001. Cohen’s *d*_Contro_&_+Inhibi_t_o_rs > 1.

**Figure S4.**
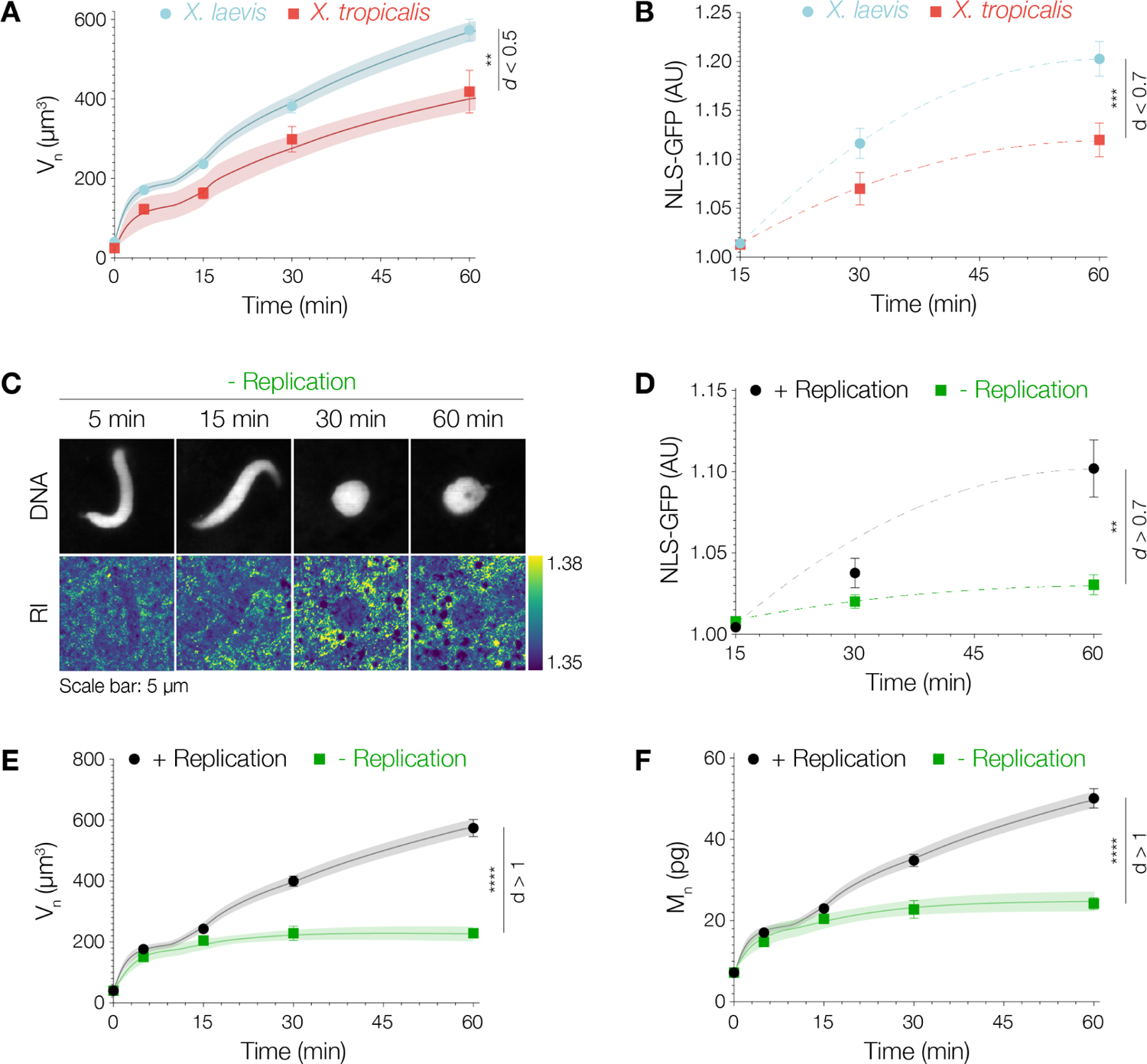
Chromatin contributes to the pressure balance both directly as a confined polymer-like chain and indirectly as a regulator of nuclear import. (A) Quantification of nuclear volume (Vn) for nuclei assembled using *X. laevis* sperm (blue) and *X. tropicalis* sperm (red). *X. tropicalis* nuclei are approx. 30% smaller than *X. laevis* nuclei. Mann-Whitney test where ** indicates p < 0.01. Cohen’s *d*_Laevis_&-r_ro_p_icalis_ < 0.5. In panels (A), (E) and (F), bold lines and shaded areas represent the mean ± SEM from the theoretical simulations for *X. laevis* and *X. tropicalis* nuclei. See Supplemental theory. (B) Quantification of NLS-GFP intensity during nuclear assembly. *Xl* nuclei are more import efficient in comparison to *Xt* nuclei. Mann-Whitney test where ** indicates p < 0.01. Cohen’s *d*_laevis_&_tro_p_s_ < 0.7. (C) Nuclear assembly in the presence of the replication inhibitor Aphidicolin. Top row shows fluorescence images of DNA stained with Hoechst-33342 and bottom row shows the RI distribution in the central slice from the ODT tomograms. Scale bar: 5 μm. Color bar on the right shows the RI distribution. (D) Replication competent nuclei (+ Replication, black circles) are more import efficient in comparison to perturbed nuclei (- Replication, green squares). Quantification of NLS-GFP intensity at different time points of nuclear assembly (n = 30 from 3 independent experiments). Symbols and bars show mean ± SEM. Mann-Whitney test where ** indicates p < 0.01. Cohen’s *d*_Laevis_&-r_ro_p_s_ > 0.7. (E) Blocking replication reduces nuclear volume. Quantification of V_n_ at different time points after the start of nuclear assembly (n = 30 from 3 independent experiments). Volumes (mean ± SEM) V_n_5_ =151 ±9 pm^3^, Vn_15 =211 ±11 pm^3^, Vn__30_ = 241 ± 25 μm^3^, and V_n_60_ = 234 ± 15 μm^3^. Volume values for + Replication data are similar to values shown in Figure 1E. Mann-Whitney test where **** indicates p < 0.001. Cohens *d*+Replication&-Replication > ^1^. (F) Blocking replication reduces nuclear dry mass. Quantification of M_n_ at different time points of nuclear assembly (n = 30 from 3 independent experiments). Dry mass (mean ± SEM) M_n_5_ = 14.8 ± 0.9 pg, M_n_15_= 20.5 ± 1.1 pg, M_n_30_= 21.7 ± 1.9 pg, and M_n_60_= 24.2 ± 1.4 pg. Dry mass values for + Replication data are similar to values shown in Figure 1E. Symbols and bars show mean ± SEM. Mann-Whitney test where **** indicates p < 0.001. Cohen’s *d*+Replcaton&-Replcaton > 1.

**Figure S5.**
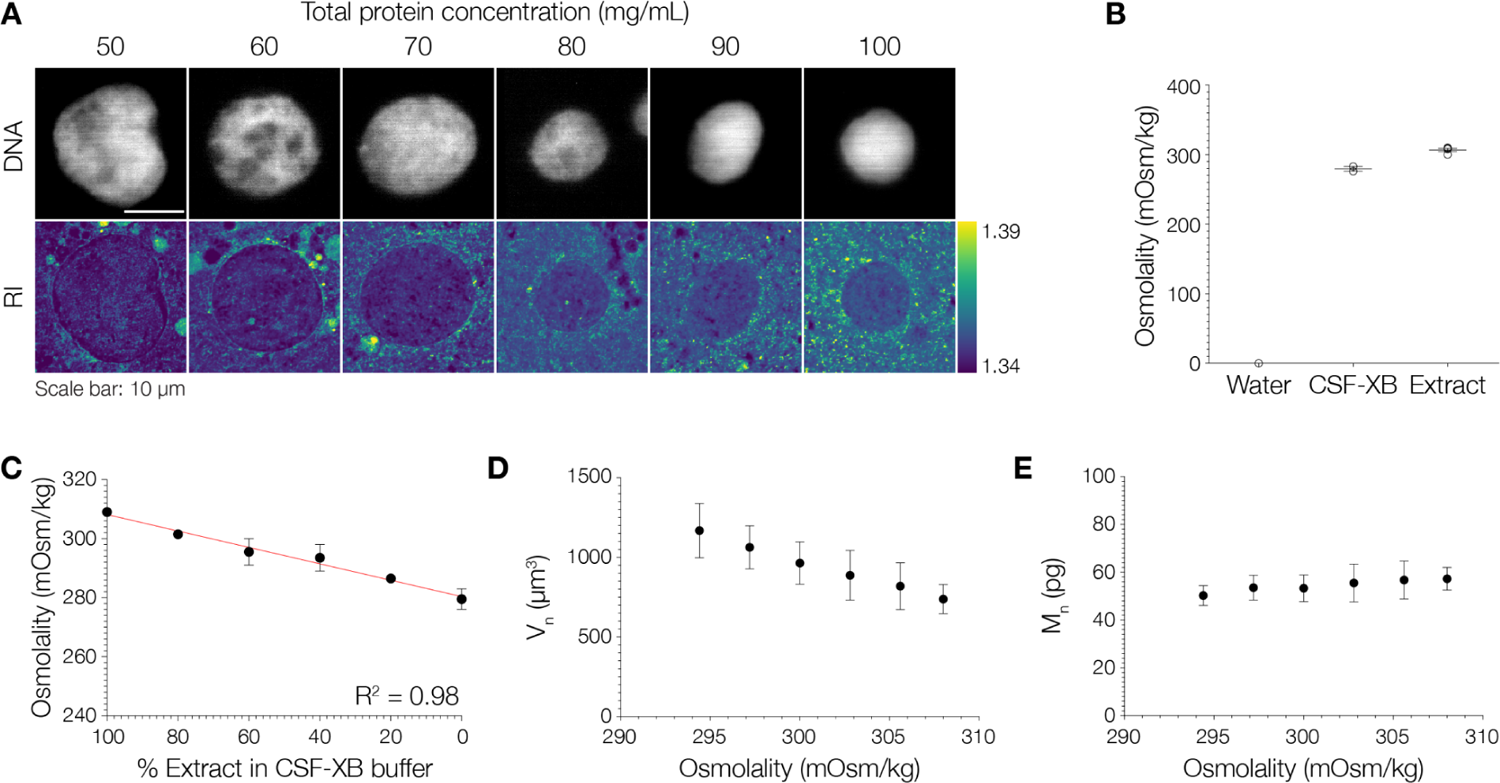
Nuclei assembled in *Xenopus* egg extract behave as near-ideal osmometers. (A) Nuclei alter their volume in response to changes in total protein density and correspondingly osmotic pressure. Representative fluorescence and RI images of nuclei assembled in *Xenopus* egg extract with reducing protein density by dilution with CSF-XB buffer. Undiluted extract has a protein concentration of 100 ± 2 mg/mL. Scale bar = 10 μm. Bar shows RI distribution range. (B) Osmolality (b) of different solutions as measured by a freezing point osmometer. b_water_ = 0 mOsm/kg (n=3), b_CSF.XB_ = 280 ± 3 mOsm/kg (n=2) and b_extract_ = 307 ± 2 mOsm/kg (n=4). Bars represent mean ± SEM. (C) Dilution of *Xenopus* egg extract in CSF-XB buffer linearly reduces osmolality of the extract (n=2 independent experiments). b_100%Extract_ = 309 ± 1 mOsm/kg, b_80%Extract_ = 301 ± 1 mOsm/kg, b_60%Extract_ = 296 ± 5 mOsm/kg, b_40%Extract_ = 294 ± 5 mOsm/kg, b_20%Extract_ = 287 ± 1 mOsm/kg and b_100%buffer_ = 280 ± 3 mOsm/kg. Circles and bars represent mean ± SEM. Red line shows a fit via simple linear regression. (D) Reconstituted nuclei in *Xenopus* egg extracts behave like near-ideal osmometers. Reducing osmolality (by altering cytoplasmic protein density via dilution with CSF-XB buffer) alters nuclear size (n=10 nuclei for each condition). V_n_294mOsm/kg_ = 1168 ± 171 pm^3^, V_n_297mOsm/kg_ = 1063 ± 135 pm^3^, V_n_300mOsm/kg_ = 964 ± 133 pm^3^, V_n_303mOsm/kg_ = 887 ± 156 pm^3^, V_n_306mOsm/kg_ = 819 ± 146 pm^3^ and V_n_309mOsm/kg_ = 738 ± 92 μm^3^. Circles and bars represent mean ± SEM. (E) Reducing osmolality does not affect nuclear dry mass (mean ± SEM, n = 10 nuclei for each condition). M_n_2g4mOsm/kg_ = ^50^ ± ^4 pg^, Mn_297mOsm/_kg_ = ^53^ ± 5 pg, M_n_300mOsm/kg_ = ^53^ ± ^6 pg^, M_n_303mOsm/kg_ = ^56^ ± 8 pg, M_n_306mOsm/kg_ = 57 ± 8 pg and M_n_30gmOSm/kg_ = 57 ± 5 pg.

**Figure S6.**
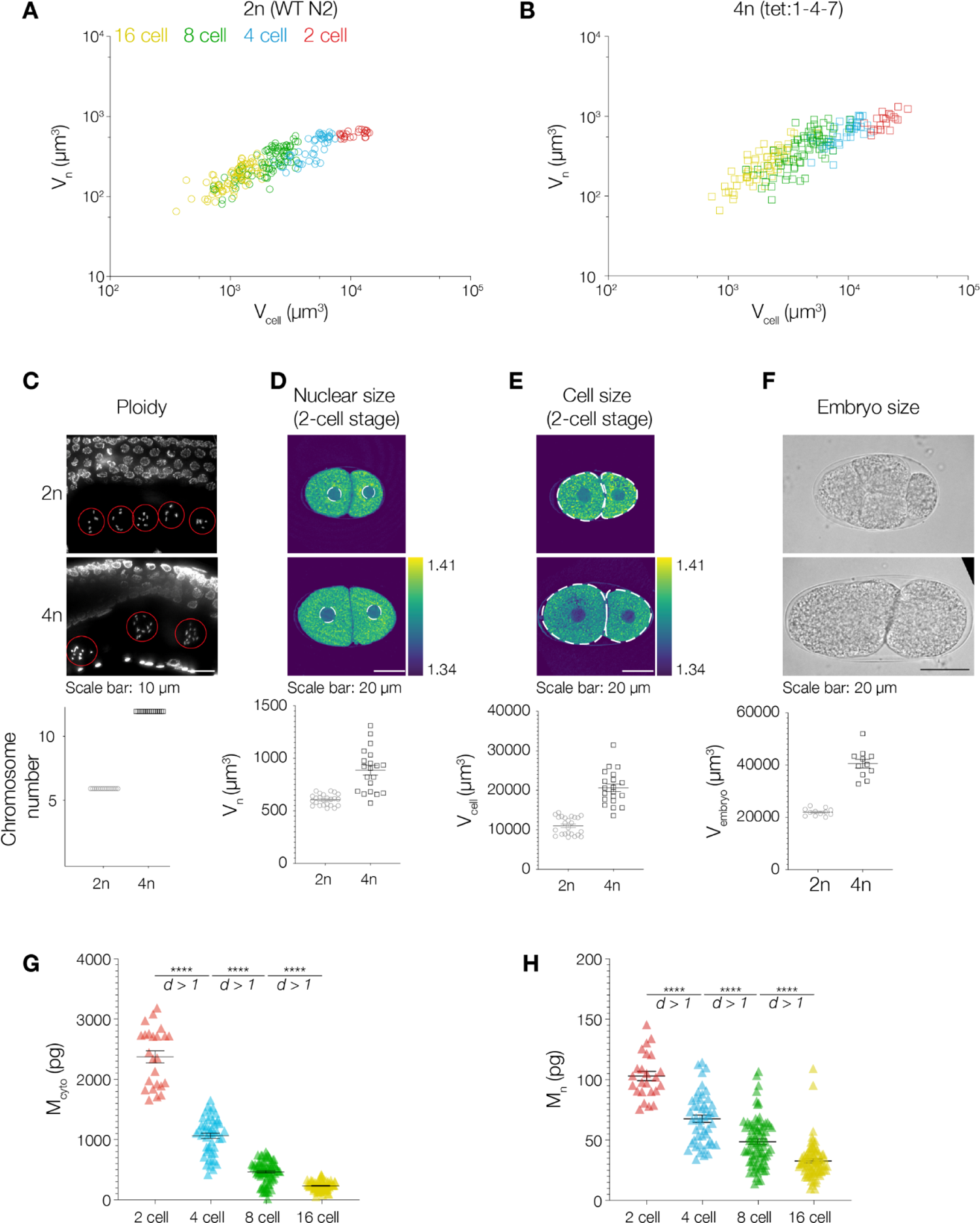
Hierarchical size scaling in *C. elegans*. (A) Double logarithmic plot showing how nuclear volume scales with cell volume in wildtype diploid *C. elegans* embryos. Red, blue, green, and yellow circles represent values from 2 cell stage (n = 24), 4 cell stage (n = 48), 8 cell stage (n = 79), and 16 cell stage (n = 85) cells, respectively. (B) Double logarithmic plot showing how nuclear volume scales with cell volume in tetraploid *C. elegans* embryos. Red, blue, green, and yellow squares represent values from 2 cell stage (n = 20), 4 cell stage (n = 44), 8 cell stage (n = 80), and 16 cell stage (n = 58) cells. (C) Representative Hoechst-33342 stained images showing the number of chromosomes in nuclei (red circles) from wildtype (top) and tetraploid (bottom) gonads. Scale bar = 10 μm. Quantification of the number of chromosomes in oocytes of 1-4-7 worms (4n, n = 15) confirmed tetraploidy for all. (D) Nuclei in tetraploid embryos have a larger volume than nuclei in wildtype embryos. Representative ODT image (central slice) from a 2-stage wildtype (n = 24) and tetraploid embryo (n = 20). Nuclei marked by white dashed lines. Scale bar = 20 μm. Color bar on the right shows the refractive index range. Quantification of nuclear volume, V_n_2n_ = 605 ± 11 pm^3^ and V_n_4n_ = 888 ± 46 μm^3^. Bars show mean ± SEM. (E) Cells in tetraploid embryos have a larger volume than cells in wildtype embryos. Representative ODT image (central slice) from a 2-cell wildtype (n = 24) and tetraploid embryo (n = 20). Cells marked by white dashed lines. Scale bar = 20 μm. Color bar on the right shows the refractive index range. Quantification of cell volume, Vc_elL2n_ = 11001 ± 454 pm^3^ and V_Gell__4_n_ = 20616 ± 976 μm^3^. Bars show mean ± SEM. (F) Tetraploid embryos (n = 11) are larger than wildtype embryos (n = 12). Brightfield images of a 4-cell stage wildtype embryo and a 2-cell stage tetraploid embryo. Scale bar = 20 μm. Quantification of embryo volume, V_emb_ry_o__2_n_ = 22023 ± 392 pm^3^ and V_embr_y_G__4_n_ = 40563 ± 976 μm^3^. Bars show mean ± SEM. (G) Cytoplasmic dry mass reduces during development in wildtype embryos. M_2Gell_ = 2373 ± 100 pg, M_4Gell_ = 1056 ± 45 pg, M_8_c_ell_ = 455 ± 21 pg and M_16_c_e_n = 222 ± 8 pg. Mann-Whitney test where **** indicates p < 0.0001. Cohen’s *d* > 1 for all comparisons. (H) Nuclear dry mass reduces during development in wildtype embryos. M_n_2_c_ell_ = 103 ± 4 pg, M_n_4_c_ell_ = 67 ± 3 pg, Mn__8cell = 48 ± 2 pg and M_n_ = 32 ± 2 pg. Mann-Whitney test where **** indicates p < 0.0001. Cohen’s *d* > 1 for all comparisons.

### Supplemental movies S1-S3

1. Supplemental movie S1: A dynamic model for nuclear assembly and growth in *Xenopus* egg extracts captures the evolution of nuclear volume (Vn) over time in both control and perturbed nuclei (- Import, - Replication, *X. tropicalis*). Video: 10.6084/m9.figshare.23668305
2. Supplemental movie S2: First embryonic divisions in a wildtype *C. elegans* embryo imaged by ODT. Scale bar = 20 μm. The RI range in the movie is 1.34-1.41. Images were acquired at 1-minute intervals. Video: 10.6084/m9.figshare.23668323
3. Supplemental movie S3: First embryonic divisions in a tetraploid *C. elegans* embryo imaged by ODT. Scale bar = 20 μm. The RI range in the movie is 1.34-1.41. Images were acquired at 1-minute intervals. Video: 10.6084/m9.figshare.23668332

## Supplemental theory

## 1. Theoretical Model of Nuclear Growth

Here, we propose a model for nuclear assembly and growth starting from sperm chromatin in *X. laevis* egg extract. It uses realistic biophysical parameters and is able to correctly reproduce the time evolution of the nuclear volume *V_n_* and dry mass *M_n_* in normal conditions and various perturbation experiments. Figure S7 provides a graphical summary of our experimental observations made in Figures 1-3 of the main text: nuclear assembly and growth can be divided into two phases, which are formally separated by the closure of the nuclear envelope. Phase 1 represents the rapid decondensation of sperm chromatin by nucleoplasmin, while phase 2 shows more gradual dynamics where nucleocytoplasmic transport of proteins and DNA replication, via osmotic and entropic pressures respectively, drive nuclear growth.

The model conceptually focuses on two aspects: I) determining the size of the nucleus based on the balance of involved pressures for the experimentally determined protein content in the nucleoplasm and cytoplasm at fixed time points, and ii) describing the dynamic nuclear dry mass as a result of nuclear transport, thus recapitulating the complete time-dependent nuclear growth.

We first detail the concept of pressure balance that is naturally applicable in phase 2 of nuclear growth, where the nuclear compartment is well defined. We discuss the idea of protein colloid osmotic pressure and chromatin entropic pressure, as well as the role of protein complexes and protein identities. By using the pressure balance concept, we can recapitulate the experimental observations on nuclear volume and mass (see 1.1). We then extend this concept to phase 1 of nuclear assembly which helps us to define the effect of the excluded volume of chromatin (1.1.4). In phase 1 and 2, chromatin experiences changes first due to decondensation and then due to replication, respectively, which we describe as a function of time (1.1.5). We next explain how the nuclear protein content evolves over time due to active nuclear import and quantify it by means of the kinetic model (1.2). By combining the concept of pressure balance with nuclear import and taking time-dependent parameters (chromatin decondensation and content) into account, we provide a complete dynamical model of nuclear growth linking both phases (1.2.2). In the concluding 1.3, 1.4, and 1.5 we explain the choice of model parameters, demonstrate the validity of the model for perturbation experiments and provide technical details of model implementation, respectively.

**Figure S7:**
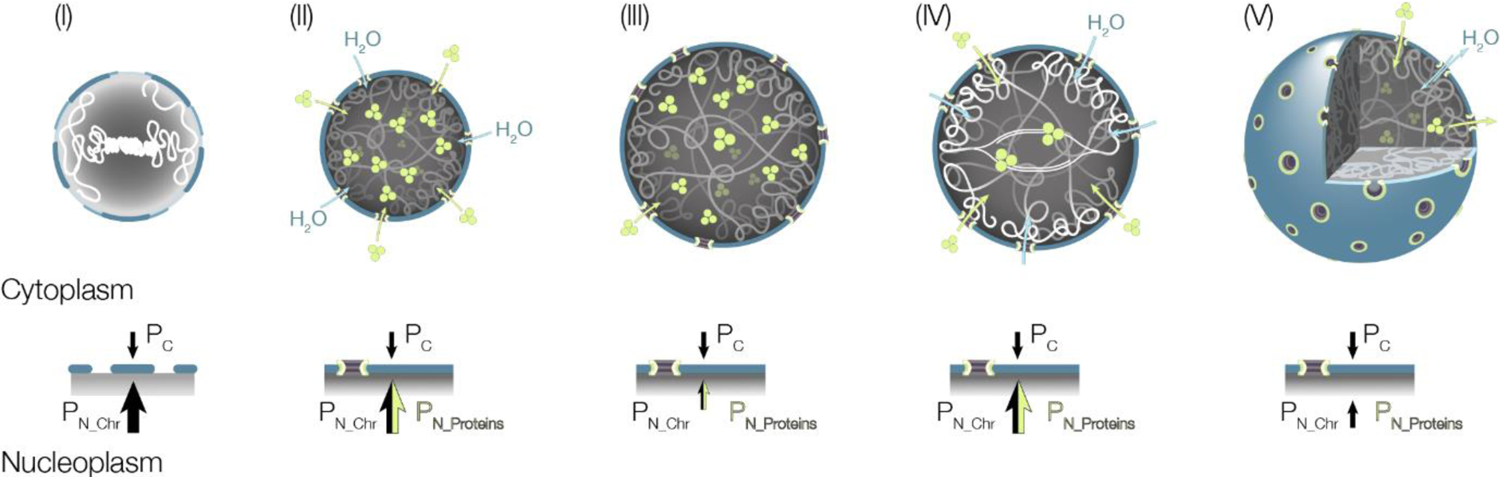
Summary of essential processes happening during nuclear growth Decondensation of sperm chromatin driven by nucleoplasmin leads to its unfolding while the nuclear envelope forms [(I), / = 0-5 min]. The fully functional nuclear envelope marks the beginning of phase 2 [(II), *t =* 10-15 min]. Specific proteins are imported and the volume of the nucleus grows (III). DNA replication starts at around 20-30 min (IV) and is finished by / = 60 (V).

### 1.1. Pressure balance

We first introduce the concept of pressure balance in phase 2, when the nuclear envelope is formed. This allows us to clearly define the nucleoplasm and cytoplasm, their respective osmotic pressures, and chromatin pressure.

#### 1.1.1. Pressure balance during phase 2

In phase 2, we assume that the nucleus possesses a fully functional nuclear envelope (NE) maintaining the compartments’ identity and integrity. Following previous work (*13, 48–50*), we consider the nuclear volume to be set by the balance of pressures present in the system. In this picture, inward and outward pressures are balanced by the NE where any imbalance results in changes of the nuclear volume via water flux across the nuclear envelope.

We assume the pressures originating from the nuclear and cytoplasmic compartments to be isotropic, which is supported by the nearly spherical nuclear shapes at large enough times *(t* > 15). In a cellular context, non-isotropic pressures can originate from interactions of the NE with the cytoskeleton which, however, has no contribution in our cell-free system. We can also neglect nuclear envelope tension as supported by Fig. S5 where we show that the *X. laevis* nucleus behaves like a near-ideal osmometer for all regimes tested in this work (as has been shown previously for the *S. pombe* nucleus in (W

Therefore, the pressure balance equation *(49)* can be written:

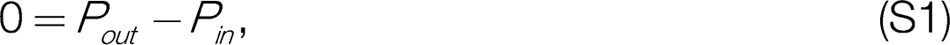

where *P_out_* denotes the outward pressures increasing nuclear volume and *P_in_*, the inward pressures decreasing nuclear volume. It has been recently proposed that the osmotic pressures generated by nuclear and cytosolic proteins, respectively *(45-50)* are the key players contributing to this balance (see Fig. 4A, 5D and H).

#### 1.1.2. Colloidal osmotic pressure

In the context of osmotic pressure, the nuclear envelope is not simply a semipermeable membrane. The NPCs act as the gateways between the nucleus and cytoplasm through which small molecules can freely diffuse, whereas molecules larger than — 30 - 40 kDa cannot (*11 i).* As a result, small macromolecules and ions despite being major osmolytes, do not contribute to the pressure difference (see, however, the discussion of the role of counterions in the main text). Larger proteins need to be specifically transported into the nucleus to establish the nuclear proteome *(51, 112-114)* (see below, 1.2.1). Therefore, to estimate the osmotic pressure, we consider the contribution of localized proteins and neglect the effects of ions and other small molecules. In this case, colloid osmotic pressure defines the source of the osmotic pressure, which are mainly proteins *(46).* We define A*P_osm_* as the difference between the outward osmotic pressures exerted by nuclear proteins and the inward osmotic pressure exerted by cytoplasmic proteins, which are determined by the Van’t Hoff law (*115*)*’*.

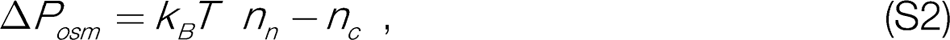

where *k_B_* is Boltzmann constant, Tis temperature, *n_nc_* denote the protein concentrations in the nucleus and cytoplasm, respectively. To estimate the protein concentration in the cytoplasm, we use the measured density *p_c_* =100 mg/mL. The cytosolic and nucleoplasmic compositions of proteins are characterized by a number average protein mass *m_c_* and *m_n_*, respectively. We also consider that proteins *in vivo* exist as parts of larger complexes, which increases the effective *m_cn_* and consequently reduces the overall protein concentration (*46, 113, 116*). Using quantitative mass spectrometry data from *X laevis*oocytes (*51*), we take *m_c_* = 155 kDa (weighted average of protein complexes) and calculate the concentration of cytoplasmic protein complexes to be *n_c_=p_c_*/m_c_ 0.65 mM. Given that macroscopic amounts of extract are present in our system, we assume that the formation of a nucleus does not deplete the extract and thus cytoplasmic quantities such as *n_c_* and *m_c_* remain constant throughout nuclear assembly and growth.

While we can directly use density measurements to calculate protein concentrations for the cytoplasm (*28*), this is different for the nucleus. Here, we need to consider the contribution of chromatin to i) the density measurements and ii) the nuclear volume available to proteins.

Chromatin has a negligible contribution to the osmotic pressure but it does contribute noticeably to nuclear mass and occupied volume. Therefore, the protein fraction of the nuclear dry mass at a given moment of time can be calculated from the total dry mass of the nucleus as *M_n_ = M_tnt_-M_rhr_.* To estimate the mass of chromatin *M_chr_,* we add up the masses of base pairs and nucleosomes:

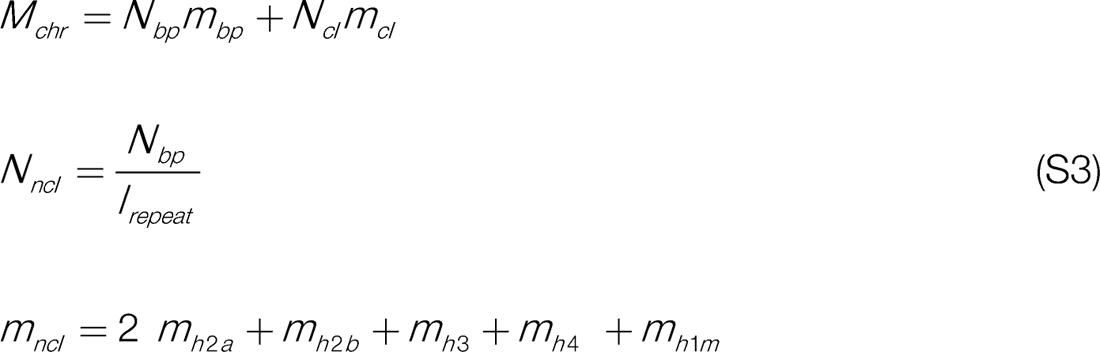

where *N_bp_* is the number of base pairs in the genome, *m_bp_* denotes the mass of a base pair (0.65 kDa (*117*)*), N_nd_* is the number of nucleosomes which can be estimated with the knowledge of the nucleosome repeat length *I_repeat_* (200 bp (*118*)). The mass of a nucleosome *m_nd_* is calculated as the added mass of the histone octamer consisting of two copies of the histones H2A, H2B, H3, and H4 each and one copy of the maternal linker histone H1M. For the *X. laevis* genome of 3.1Gbp this results in *M_chr_* = 6.9 pg (and double of that after replication). Considering the weighted average of protein complexes in the nucleus leading to *m_n_* = 131 kDa (57), we can calculate the protein concentration in the nucleus as *n_n_=M_n_/ m_n_V_n_*.

If we now use the above equations and measurements of dry mass to calculate the difference of osmotic pressures in (S2), we would arrive at *&P_osm_* ∼ −330 Pa (at *t* = 60 min) meaning that the nucleus would rather shrink than grow at this negative pressure difference. By repeating the same calculation of the predicted volumes of the nuclei based on the condition of osmotic pressure balance for all time points, we arrive at the first set of points in Figs. 4C and D (empty squares) that noticeably underestimate the volume. We suggest that this discrepancy can be corrected by considering chromatin pressure.

#### 1.1.3. Chromatin pressure

Next, we consider the outward pressure that chromatin can exert. As chromatin decondenses, it attains its thermodynamically preferred conformation quantified by the radius of gyration *R (119).* Confinement by the NE, however, results in a nuclear radius smaller than the gyration radius and thus leads to an outward pressure exerted by chromatin on the NE. To quantify the relevance of this effect, we describe chromatin as one semi-flexible, self-avoiding polymer chain, which was successful in quantitative description of DNA in various settings *(120-122).* For such a polymer confined in a sphere, the chromatin pressure is defined by *(49, 123):*

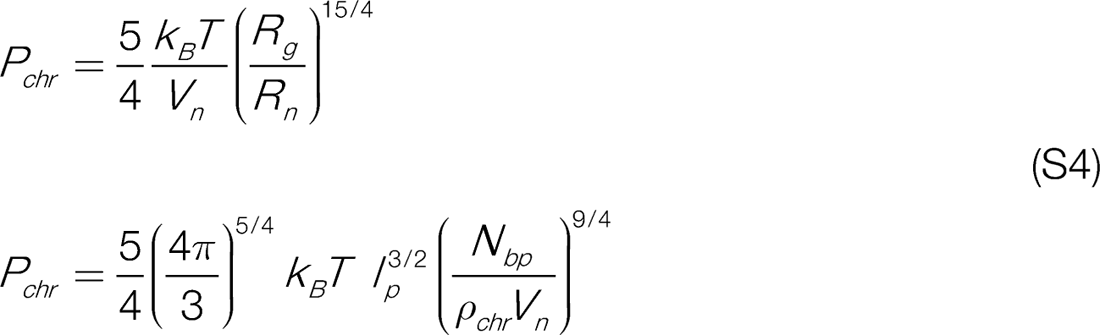

The radius of gyration can be estimated as R_g=_ I_p_^3/2^L_c_^3/5^ (52) with the contour length of polymer *L_c_* and its persistence length *I*. We can estimate *L_c_* with the knowledge of the total number of base pairs in the genome *N_bp_* and the linear density of chromatin *p_chr_* that describes the number of base pairs per unit length via *L_c_ = N_bp_/ p_chr_.* For the persistence length, we find literature values in the range of 100-200 nm (124), and for the linear density it is 20-90 bp/nm *(118, 124, 125).* For the first preliminary estimate, we take a linear density value of 40 bp/nm (*118, 124)* and a persistence length of 150 nm, roughly in the middle of literature range.

If we consider the nucleus at *t* = 60 minutes, one round of DNA replication has been completed doubling the number of base pairs to *N_bp_* =6.2 Gbp and resulting in *R_g_* ∼ 600 pm. The average volume of the nucleus at this time is Ri600 pm^3^, corresponding to a nuclear radius *R_n_* 5 pm, which is significantly smaller than *R_g_*. Thus, chromatin will contribute to the outward pressure with *P_chr_* 540 Pa. This value is on the order of magnitude to compensate for the above calculated osmotic pressure difference (see Figs. 40 and D). The osmotic pressure due to localized proteins tries to shrink the nucleus as the cytoplasm is more concentrated than the nucleus, whereas the chromatin pressure expands the nuclear volume.

It is worth noting that in another study *(49),* the estimates for chromatin vs osmotic pressures led to almost two orders of magnitude difference, which ultimately led the authors to neglect the contribution of chromatin. The reason for such a difference is twofold. First, the individual proteins (i.e. polypeptide length) were used to calculate concentrations (not the complexes) and the absolute values of pressures (not their difference) were considered, which together led to a much higher estimate for the osmotic pressure term. Second, chromatin parameters in *(49)* rely on values for a unicellular system, yeast. We adopted the values reported for somatic systems with lower chromatin compaction thus leading to higher chromatin pressure in our case. Therefore, our estimates follow the conceptual logic of *(49)* but use parameters more pertinent to the system considered in this work and, taken together, support the commensurable effects of colloid osmotic pressure difference of cytosolic/nuclear compartments and entropic chromatin pressure.

We can now write the simple pressure balance (which for now ignores the effect of the excluded volume of chromatin, see below) starting from equation (S1) and substituting the osmotic pressure (S2) and entropic chromatin pressure (S4):

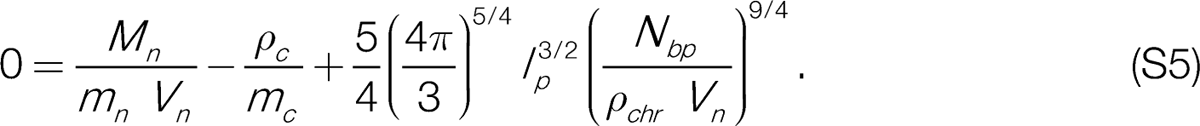

Equation (S5) was used for the chromatin-pressure-corrected set of nuclear volumes plotted in Figs. 4C and D of the main text (empty diamonds), which now much better match our experimental data.

In the subsection 1.1.5 below, we will use the pressure balance and experimental measurements of dry mass to calculate the respective chromatin characteristics at different time points. However, before doing that we have a quick look at how the concept of pressure balance can be useful in phase 1, when there are no well-defined compartments yet.

#### 1.1.4. Excluded volume of chromatin

In phase 1, the nuclear envelope is still forming and thus we can not yet define the nucleus as a separate compartment. Therefore, we formally define nuclear volume as the volume encompassing the chromatin. From the measured dry mass of the nucleus we can calculate the mass of nuclear proteins at *t* = 5 min (we should note that the first time point of *t* = 0 measured for sperm chromatin is just in a buffer solution). Proteins in the available nuclear volume should have equal osmotic pressure to the cytosolic proteins and thus equal concentrations:

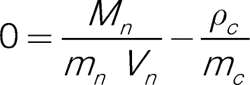

Using *m_n_* (for simplicity we assume the same nuclear identity of proteins at all time points) and the corresponding protein mass *M_n_*, we can calculate protein number. Using the above equation, we estimate the effective volume available to proteins which is *Vf* ∼ 120 pm^3^ (at *t* = 5 min). This volume is lower than the measured volume *V_n_* 170 pm^3^ of the nucleus. This suggests that the volume available to proteins is reduced by the space occupied by chromatin. We can then define the effective excluded volume of the chromatin as *V_chr_=V_n_-*V_n_^eff^, resulting in *V_chr_* 50 pm^3^ at *t* = 5. This result is very close to the estimated minimal value of the physical chromatin volume calculated as the sum of the DNA and nucleosomes’ volumes *(126)* for the replicated *X. laevis*genome ∼ 24 pm^3^, see Table S1 for the exact parameter values.

We assume that for later times the excluded volume of chromatin is not changing till the onset of replication and doubles thereafter. Importantly, the excluded volume of the chromatin will limit the space available for proteins with consequences for protein colloid osmotic pressure in the nucleus:

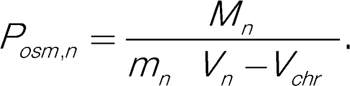

#### 1.1.5. Chromatin decondensation and replication

During the course of our observations, chromatin condensation and content are changing. In phase 1, it is the biochemical and steric change leading to its decondensation, while in phase 2, it further relaxes to its equilibrium shape and then proceeds through replication. For simplicity of analytical treatment, we describe chromatin in all stages as a self-avoiding semi flexible polymer with parameters of number of basepairs *N_bp_*, persistence length *l_p_*, and linear density *p_chr_*. The former two can be combined in a parameter *k = l^2^_p_^/5^p∼_b_’^5^* that determines the gyration radius of the polymer via *R_g_= k* and thus its pressure when in confinement. With this definition and with the excluded volume correction introduced in section 1.1.4, the final pressure balance equation is given by:

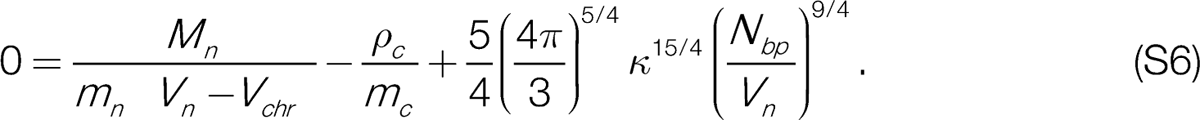

Generally, it is difficult to measure chromatin properties directly. We can, however, use our experimental data to determine *n(t)*. In phase 1, we obtain *k* by simply equating the measured nuclear volume to the volume of the sphere with *R = R_g_* (Note, however, that it is a strong approximation as shapes of the nuclei at early stages are not spherical yet). In phase 2, we can get the value of from the pressure balance equation (S6) where the osmotic pressures difference is equilibrated by the chromatin pressure with a given k and *N_bp_* (note that *N_bp_* doubles from around 30 to 60 min due to replication). Thus, calculated plotted as a function of time in Fig. S8 reveals the change of two orders of magnitude between phases 1 and 2. It is important to remember, however, that in phase 1, sperm chromatin is initially highly condensed and thus its description via a self-avoiding polymer with the parameter *n* during decondensation is at best phenomenological. However, for times >15 min it should converge to biologically realistic parameters, which it does (as shown by the red-star symbol in the plot calculated for /_p_ = 150 nm and *p_chr_* = 40 bp/nm). We next approximate with two fitting functions: i) Michaelis-Menten growth in phase 1, and ii) sigmoidal (exponential) relaxation in phase 2 (see Fig. S8 caption for details), thus containing a minimum of free parameters. We set (without loss of generality) the boundary of the two phases to *t* = 10 min (blue symbol), where we also see the onset of nuclear transport in experiments.

So far, we rationalized the relationship between the measured volume of the nucleus, the mass of nuclear proteins, and chromatin at different time points via the pressure balance. We next wanted to understand the time dynamics of nuclear growth.

**Figure S8:**
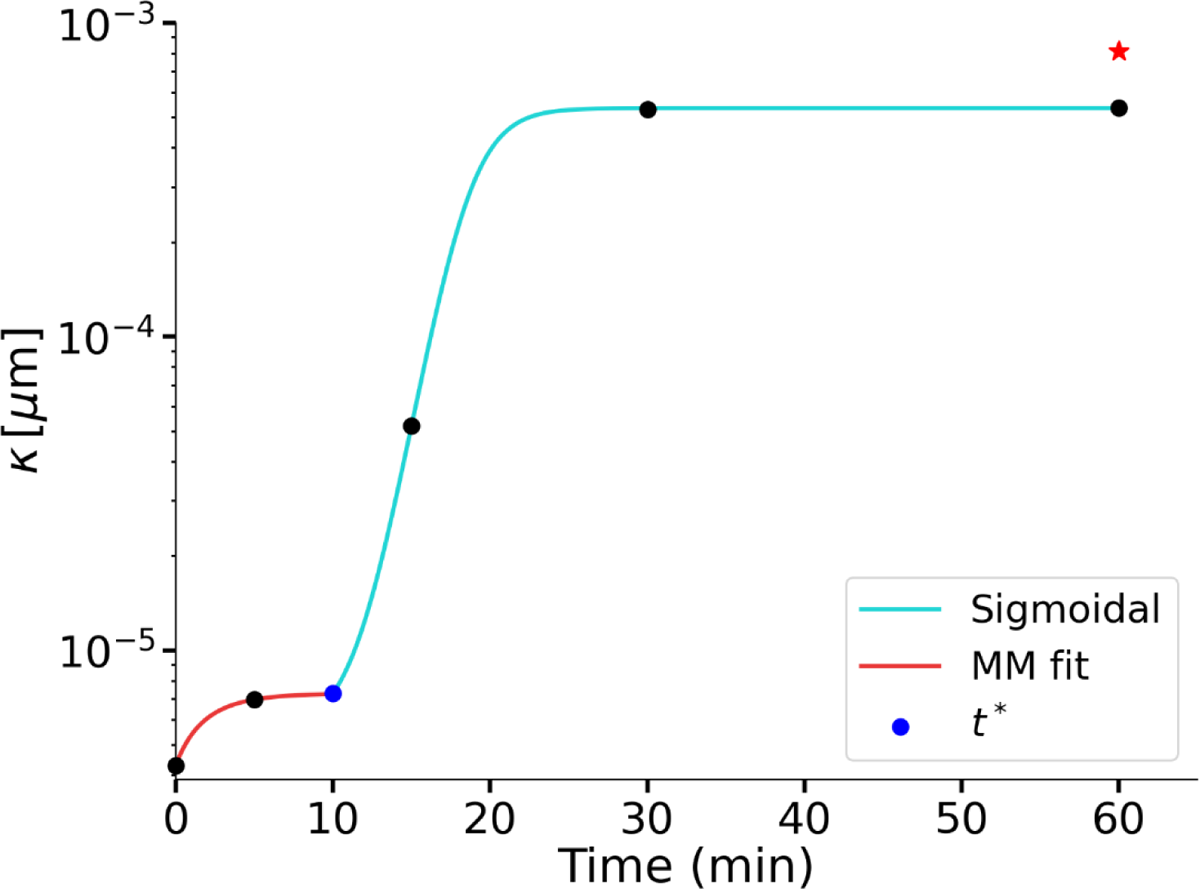
Quantifying parameter *k* from the pressure balance. Values of *k* are plotted in phase 1 *(t* <10min) from equating the measured volume of the space occupied by chromatin to a sphere with a gyration radius as function of k (see the text), in phase 2 (/> 10min) from the pressure balance equation (S6). The data points are then fitted with a Michaelis-Menten kinetics in phase 1 and simple sigmoidal behavior in phase 2. The fit indicates that once chromatin fully decondenses at around 20 min, k and therefore its mechanical properties do no longer change. The star-symbol represents the k values calculated for literature values of persistence length !_p_ = 150 nm and compaction ratio *p_chr_* = 40 bp/nm. The blue symbol denotes the transition time between the two phases *t* =10 min.

### 1.2. Dynamic model

Our next step is to expand the model enabling us to predict nuclear volume *V_n_* and protein content *N_n_* as a function of time. In phase 1, the volume of the nucleus just follows the decondensation of chromatin, and the protein mass follows cytosolic concentration in the volume of the nucleus reduced by the excluded volume of the chromatin. In phase 2, dynamics are richer. If the pressures in equation (S6) are not balanced, the nucleus adjusts its volume via water flux until the balance is reached, which can be described by the following equation:

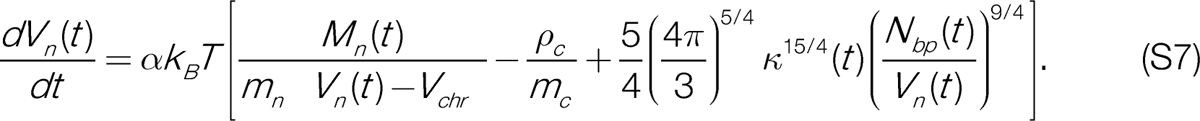

where *a* is the phenomenological constant describing hydraulic permeability of the nuclear envelope (it will not affect calculations under the assumption of fast water equilibration *(48, 127, 128)).* During phase 2, active nucleocytoplasmic transport establishes the nuclear proteome. This causes changes in the number of protein complexes in the nucleus *N_n_= M_n_ / m_n_,* which in turn affects the osmotic pressure in equation (S7). Thus, our model requires a mathematical description of nucleocytoplasmic transport. This transport system has been extensively studied over the years and several models have been proposed, ranging from conceptual to rather detailed *(62, 112, 129)*.

#### 1.2.1. Kinetic theory of nuclear import

Nuclear pore complexes connect the nuclear and cytoplasmic compartments and only small macromolecules <30-40 kDa (*111*) are able to freely translocate through them. Larger proteins need to be selectively transported through the pores by nuclear transport receptors, such as importins. Importins bind to cargo proteins via their nuclear localization signal (NLS). The resulting cargo-importin complex is able to passively shuttle between the nuclear and cytoplasmic compartments, enabled by specific interactions of the importin with NPCs. Accumulation of cargo in the nucleus is achieved by the asymmetric distribution of the GTPase Ran: the nucleus is enriched with GTP-bound Ran (RanGTP), while the cytoplasm is enriched with GDP-bound Ran (RanGDP). RanGTP has a high binding affinity to importins, which along with its abundance in the nucleus leads to the formation of RanGTP-importin complexes, thus dissolving the cargo-importin complexes and delivering cargo in the nucleus (*129, 130*). This is how specific proteins accumulate in the nucleus. The asymmetric distribution of RanGTP is achieved by a dedicated molecular machinery. The GTPase-activating protein RanGAP, which is localized to the cytoplasmic side of the NE, hydrolyses the RanGTP of outgoing RanGTP- importin complexes, thus dissolving the complex and producing free RanGDP and importin as a result. Another transport receptor NTF2 binds to RanGDP in the cytoplasm, forming a complex that shuttles Ran back to the nucleus. Once the complex enters the nucleus, the guanine-nucleotide exchange factor RCC1 (or RanGEF), which is localized to chromatin (*131, 132*), converts RanGDP into RanGTP. This dissolves the complex and liberates RanGTP in the nucleus. In summary, the accumulation of specific proteins in the nucleus is achieved by the action of nuclear transport receptors such as importin and the asymmetric distribution of RanGDP/RanGTP, which is maintained by the localization of RanGAP to the outside and RanGEF to the inside of the nucleus. Since this process consumes energy by hydrolysis of RanGTP to RanGDP, it is also referred to as a directed active transport. Throughout the years, multiple biological and biophysical models of varying complexity have been proposed by various authors *(63, 129, 133-137).* For the reconstituted nuclei of *X. laevis,* Kopito and Elbaum *(62)* put forward an experimental demonstration and proposed a simple kinetic theory of this process (see also (*112, 129*)). Here, we build up on the model of (*112*) by explicitly linking the transport equation to the chromatin-dependent RanGEF concentration and consider it in the context of dynamically expanding nuclei as governed by the pressure balance. One of the findings of *(63)* is that accumulation of cargo in the nucleus of fixed size can be described by an effective first order kinetics equation. Concentration of the NLS cargos follows a simple differential equation:

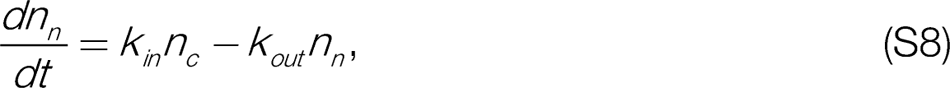

where *k_in_, k_out_* respectively denote the effective forward and backward rates of the first-order kinetics and *n_cn_* are the concentration of NLS cargo in the two compartments. For clarity of argument, we assume that *n* represents an effective single species NLS protein^1^. We argue that these effective kinetics, driven by the Ran gradient, will depend on RCC1 (RanGEF) concentration, which in turn correlates with chromatin content (*138).* To show how the effective dynamics depend on RanGEF concentration, we reproduce the core reactions of the import cycle (see (*112))*.

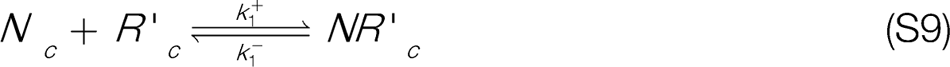

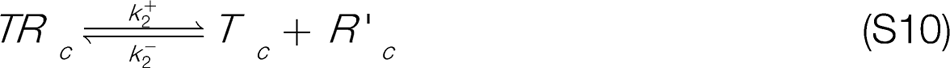

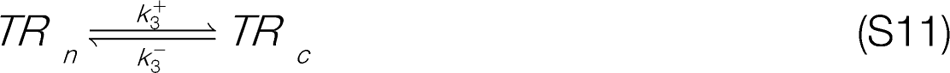

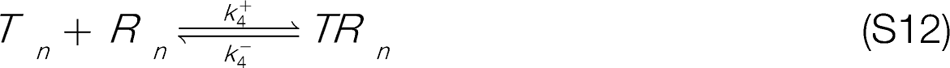

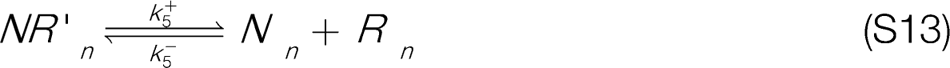

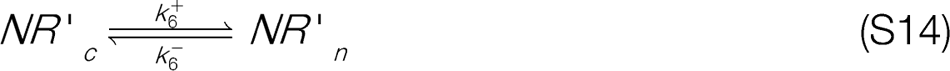

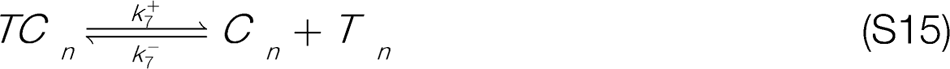

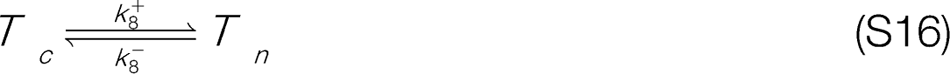

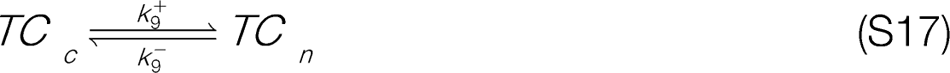

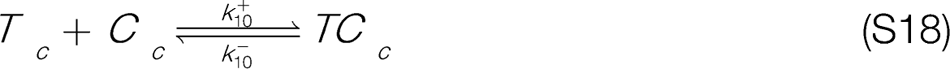

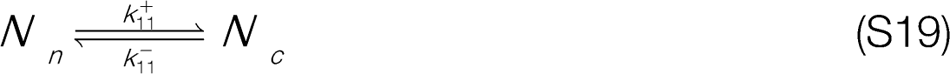

where the subindexes *n, c* denote nuclear and cytoplasmic quantities, respectively. The abbreviations we used stand for N : NTF2, R : RanGTP, R’: RanGDP, T : Importin, C : Cargo Protein. We can classify these reactions in three types: binding/ unbinding in the cytoplasm (reactions (S9), (S10), (S18)), binding/ unbinding in the nucleus (reactions (S12), (S13), (S15)) and passive diffusion through the NPCs (reactions (S11), (S14), (S16), (S17), (S19)). For this set of reactions, the corresponding set of 14 standard chemical rate equations can be written down, for which we refer to the original work *(112)*.

*We* note that reaction (S13) is where the nuclear NTF2-RanGDP is dissociated and RanGEF exchanges RanGDP for RanGTP. Therefore, the influence of RanGEF is captured in the forward rate *k_5_^+^* leading to the assumption that *k_5_^+^ RanGEF _n_*. In a similar fashion, we can make the assumption *k_5_^+^* oc *RanGAP_c_*.

By considering the steady state of the full system (S9)-(S19) and comparing it with the equation of the effective first order kinetics, after considerable algebra (of. (*112))* we can derive expressions for *k_in_* and *k_out_* in terms of the parameters of the detailed transport model:

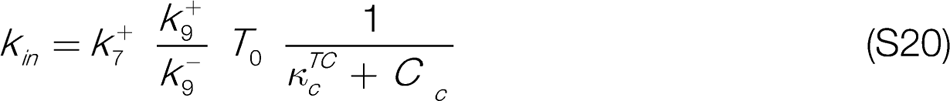

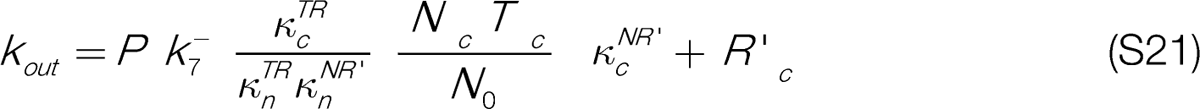

where:

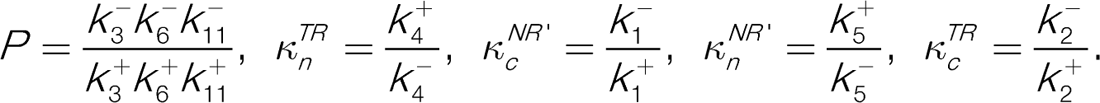

We can see that *K_in_* only depends on cytoplasmic quantities and can be indeed assumed to be constant in our model under the assumption that nuclear assembly does not affect the cytoplasmic composition. Importantly, for *k_out_* we get:

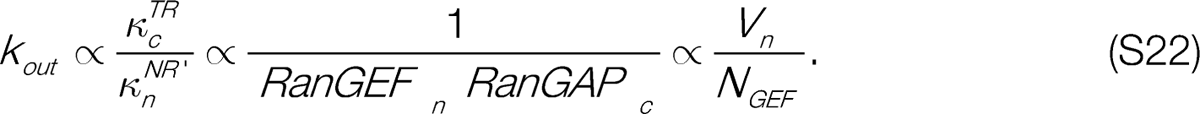

Therefore, the dependency of *k_out_* on RanGEF concentration brings in a dependency of import efficiency on the nuclear volume *V_n_*, and the amount of RanGEF in the nucleus *N_GEF_*. We assume that the amount of RanGEF in the nucleus is correlated with the amount of chromatin and would increase with replication of chromatin. This means that replication will decrease the value of *k_out_* because the value of *N_GEF_* increases. As we do not know the exact value of *N_GEF_* and how it changes during replication, we assume that replication reduces the value of *k_out_* by a phenomenological factor *v <* 1.

After using the model *(62, 112)* to establish the dependence of the effective first order kinetics of transport on the concentration of RanGEF for the steady-state volume, we are ready to push it to the dynamical regime where volume changes over time.

#### 1.2.2. Time-dependent nuclear volume dynamics

In our experimental setup, the nuclear volume *V_n_(t)* is a variable while the cytoplasmic concentration of any protein (including our effective nuclear protein) is constant, i.e. *n_c_ = const^2^.* Thus, for the time derivative of the effective nuclear protein concentration we have:

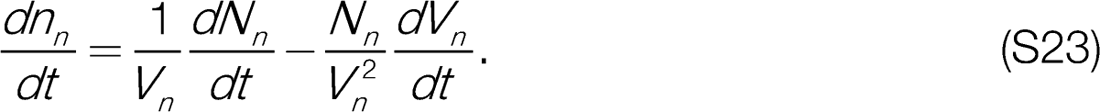

Concentration of nuclear proteins changes due to the number of protein complexes *N_n_* shuttling through the NE, but also due to the changes in its volume *V_n_.* Thus, using the first order kinetics equation (S8), we can write the time derivative for the number of nuclear proteins:

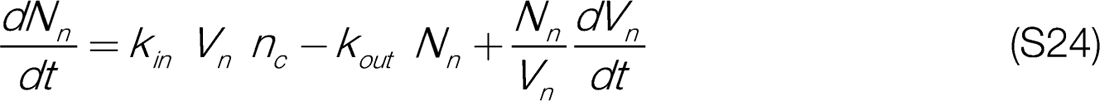

For simplicity, we can assume that water equilibration happens much faster than the characteristic time scale of nuclear import and therefore we can set *dV_n_ / dt* 0 *(48, 127, 128).* With all the considerations explained above, we arrive at the final set of equations for our model, where the nuclear volume changes in a quasisteady-state fashion. In particular, we include the RanGEF dependence of *k_out_* by means of the substitution *k_out_->k’_out_ V_n_(t) / v(t),* where *k’_out_= const* and *v(t)* reflects the maintenance of import by chromatin content:

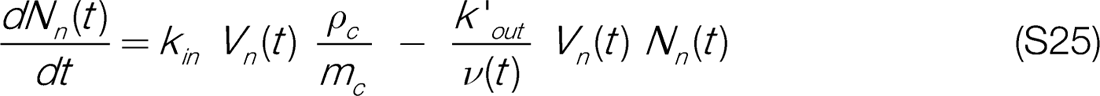

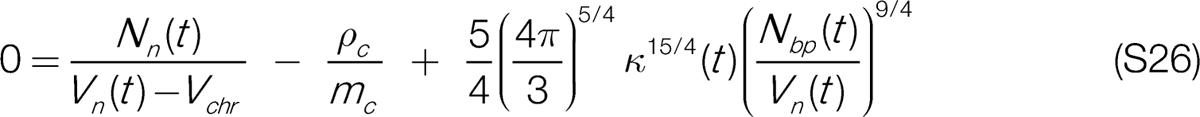

Taken together, the number of nuclear proteins changes in time following Eq.(S25) describing nuclear import, whereas the volume responds as determined by solving Eq. (S26) of pressure balance for *V_n_.* Note that *N_b_(t)* and the factor *v(t)* are time-dependent because we are incorporating chromatin replication, and characterizes chromatin decondensation (see 1.1.5 above and Fig. S8, we should note that our data suggests that replication does not effect *k* of the chromatin). The effect of replication is captured by doubling the value of *N_b_* in a linear fashion:

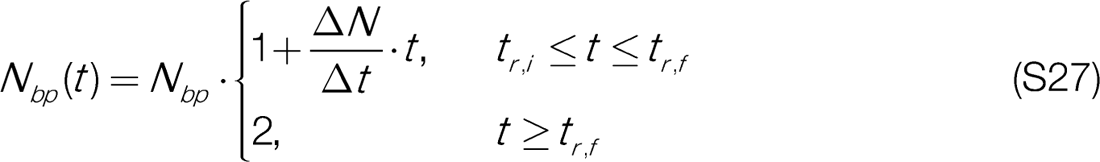

where *t_r,i_* denotes the time at which DNA replication starts, *t_rf_* the time at which DNA replication ends, *LN* = 1 and *Lt = t_rf_-t_rj_*. The value of *v(t)* changes similarly:

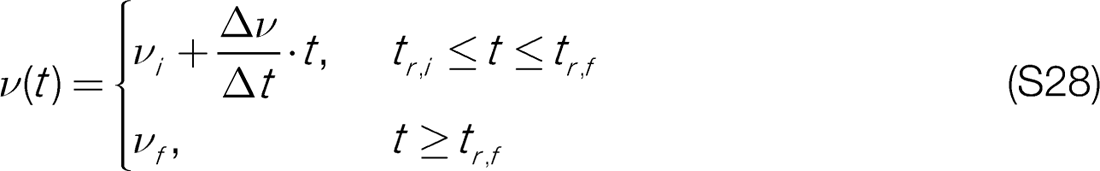

where *v_i_* = 1 and the value of *v_f_* is considered to be a model parameter, which leaves *Lv = v_f_-*1 and *Lt = t_rf_ - t_ri_*. As we will proceed to show next, the model relies on a very limited number of free parameters with most of them taken from literature or directly recovered from our experiments.

### 1.3. Parameters

We first use the control conditions to set most of the model parameters. We then show how selected parameters need to be adjusted for the respective experimental perturbations. The final time point of our experimental measurements is 60 minutes after one round of DNA replication. Therefore, we assume the final experimental time point to reflect the steady-state of the system although nuclei keep growing *(42).* This means that we know the experimental values of *N^s^_n_^s^* and *V™*. The value of is the mean volume measured at 60 minutes. The value of *N^s^_n_^s^* can be obtained with the ratio of the mean protein mass *M_n_* at 60 minutes and the number average protein mass in the nucleus *m_n_*. With this knowledge, we can estimate the value of the ratio *k’_out_! k_in_* by considering the steady-state of the transport kinetics, equation (S25):

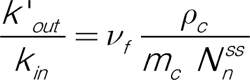

We take *k_in_* as a model parameter and can determine the value of *k’_out_* with the equation above. All in all, this leaves us with just three model parameters: *k_!n_, n(t)* and *v_f_*. All other parameters are either known from literature or directly calculated from measurements in this work. All of the biological parameter values used for the estimates in this text and in the simulations are summarized in Table S1. Additionally, the relevant model parameter values are provided in Table S2.

**Table S1:** Biological quantities and their respective values relevant to the model and calculations performed here.

| Quantity | Value | Unit | Source |
| --- | --- | --- | --- |
| Cytoplasmic density | $100 \pm 2$ | mg/mL | Our measurements |
| Nuclear density (60 min) | $91.4 \pm 0.8$ | mg/mL | Our measurements |
| Nuclear volume (60 min) | $570 \pm 30$ | $\mu\text{m}^3$ | Our measurements |
| Nuclear dry mass (60 min) | $51 \pm 2$ | pg | Our measurements |
| Cytoplasmic osmolality | $307 \pm 2$ | mOsm/kg | Our measurements |
| Protein concentration in cytoplasm (XI) | $0.65 \pm 0.02$ | mM | Our measurements |
| Protein concentration in nucleus | $0.51 \pm 0.02$ | mM | Our measurements |
| Cytoplasmic osmotic pressure (Complexes) | $1610 \pm 50$ | Pa | Our measurements |
| Nuclear osmotic pressure (Complexes, XI) | $1280 \pm 40$ | Pa | Our measurements |
| Nuclear osmotic pressure (Complexes, Xt) | $1380 \pm 50$ | Pa | Our measurements |
| Chromatin pressure (XI) | $540 \pm 60$ | Pa | Our measurements |
| Chromatin pressure (Xt) | $280 \pm 70$ | Pa | Our measurements |
| Number avg. native weight of nuclear proteins | 131 | kDa | (51) |
| Number avg. native weight of cytoplasmic proteins | 155 | kDa | (51) |
| Genome length (XI) | 3.1 | Gbp | (139) |

| Continuation of Table S1 |  |  |  |
| --- | --- | --- | --- |
| Quantity | Value | Unit | Source |
| Genome length (Xt) | 1.7 | Gbp | (139) |
| Chromatin persistence length | 100 – 200 | nm | (124) |
| Chromatin linear density | 20 – 90 | bp/nm | (118, 124, 125) |
| Nucleosome repeat length | 200 | bp | (118) |
| Avg. mass of basepair | 0.65 | kDa | (117) |
| Mass of H2.A (XI) | 14 | kDa | (140) |
| Mass of H2.B (XI) | 14 | kDa | (141) |
| Mass of H3 (XI) | 15 | kDa | (142) |
| Mass of H4 (XI) | 11 | kDa | (143) |
| Mass of B4 linker histone (XI) | 29 | kDa | (144) |
| Linear density of DNA double helix | 2.9 | bp/nm | (118) |
| Thickness of DNA double helix | 2 | nm | (145) |
| Radius of histone octamer | 5.5 | nm | (145) |
| Length of histone octamer | 66 | nm | (145) |

### 1.4. Perturbation experiments

To gain a better understanding of the mechanistic processes determining nuclear volume and density, we perturbed our experimental system in various ways. Our goal in this section is not only to fit our theory to the experimental data for the various perturbations through parameter fine tuning, but we also aim to make the parameter changes consistent with the respective perturbation.

**Table S2:** Parameter values obtained from the simulation fits for different experiments: nuclei grown under control conditions (Ctrl), under import inhibition (-Imp.), with *Xt*sperm chromatin (Xt), and finally under replication inhibition (-Rep.). See 1.5.3 for details on how the errors were obtained.

| | $k_{in}$<br>$\left[\frac{10^{-3}}{s}\right]$ | $k'_{out}$<br>$\left[\frac{10^{-6}}{s \mu m^3}\right]$ | $m_n$<br>$kDa$ | $m_c$<br>$kDa$ | $\rho_c$<br>$\left[\frac{mg}{mL}\right]$ | $\nu_f$ | $N_{bp}$<br>$Gbp$ | $V_{chr}$<br>$[\mu m^3]$ | $\kappa_{60}$<br>$[10^{-4} \mu m]$ |
| --- | --- | --- | --- | --- | --- | --- | --- | --- | --- |
| **Ctrl** | $1.43 \pm 0.07$ | $3.18 \pm 0.19$ | 131 | 155 | 100 | 1.15 | 3.1 | $48 \pm 4$ | $5.2 \pm 0.6$ |
| **-Imp.** | $1.37 \pm 0.07$ | $5.33 \pm 0.17$ | 131 | 155 | 100 | 1.15 | 3.1 | $40 \pm 9$ | $2.5 \pm 0.5$ |
| **Xt** | 1.5 | $4.20 \pm 0.17$ | 131 | 155 | 100 | 1.15 | 1.7 | $16 \pm 6$ | $6.5 \pm 1.4$ |
| **-Rep.** | $1.07 \pm 0.03$ | $6 \pm 1$ | 150 | 155 | 100 | 1 | 3.1 | $48 \pm 3$ | $2.5 \pm 0.5$ |

#### 1.4.1. Nuclear import inhibition

We start with describing the nuclear import inhibition experiments. To perturb nuclear import, an inhibitor cocktail of Ivermectin and Pitstop-2 was used. Ivermectin is a broad­spectrum Importin *a! (3* inhibitor *(40)* and Pitstop-2 is a Clathrin inhibitor that compromises nuclear permeability by modifying NPC structure (47). Upon import inhibition, the accumulation of NLS-GFP was greatly reduced (see Fig. 3B). Further, there was a reduction of nuclear volume and dry mass in comparison to control nuclei (see Figs. 3o, 3E). Importantly, the density of reconstituted nuclei did not go below the cytoplasmic density of 100±2 mg/mL (see Fig. 3D). To reproduce these conditions in our model, we reduce the influx and outflux rates *k_in_* and *k_out_.* In addition, the DNA fluorescence measurements suggest that our import perturbation also inhibits DNA replication and thus, we keep *N_bp_* constant and set *v* to 1. The inhibition of import can be expected to change the identity of the nuclear proteome, which is represented by the number average molecular weight of protein complexes *m_n_.* If import is inhibited, we expect the proteome of the perturbed nuclei to remain more similar to the cytoplasmic proteome. Therefore, we can assume *m_n_* to take values higher than in control nuclei and closer to *m_c_.* An important experimental observation in import-inhibited nuclei, is that their density is equal to that of the cytoplasm. Together this would lead to very similar osmotic pressures in the cytoplasm and nucleus. This suggests that chromatin pressure is negligible w.r.t. protein colloid osmotic pressure, which is reflected in our model by the fitted *n* values which are lower than in the control (see Table S2). Taken together, with a negligible chromatin pressure and *m_n_* being similar to *m_c_,* our theory is able to reproduce a smaller nucleus with a density almost identical to the cytoplasmic density (see Fig. S9). The import inhibition treatment has a combined effect on transport and replication. We can, however, next look at the direct effects of replication inhibition directly.

**Figure S9:**
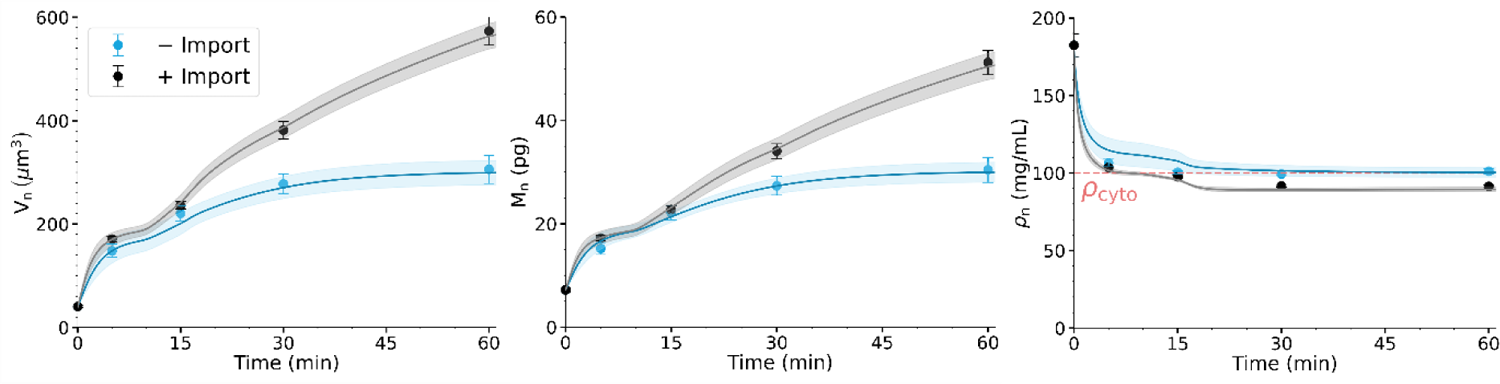
Nuclear volume, dry mass and density as a function of time for nuclei with inhibited nuclear import compared to control condition. Dots represent experimental data and solid lines are the simulation results. Error bars and shaded areas are respectively showing the SEM from experiments and simulations. Respective parameters can be found in Table S2 (-Imp. condition).

#### 1.4.2. Replication inhibition

To perturb replication, we used the DNA polymerase *a* inhibitor Aphidicolin *[146).* Aphidicolin treated nuclei were significantly smaller than control nuclei and had a lower dry mass but similar density (see Figs. S4E, F andSIO). In addition, NLS-GFP accumulation was reduced in these nuclei (see Fig. S4D).

In this condition, we keep *N_bp_* constant throughout the whole simulation as well as *v* with a value of 1, in order to mimic replication inhibition. The reduced NLS-GFP accumulation observed in this perturbation indicates that transport is also affected. This not only suggests that chromatin and transport are linked, but also allows us to change the rates *k_in_* and *k’_out_* to adjust the steady-state dry mass. As we already argued in the case of import inhibition, we can change the value of *m_n_* under the assumption that the nuclear proteome is changed due to the perturbation. Finally, given that this perturbation acts on DNA, it is reasonable to expect that chromatin properties are also going to change *(14ty,* which is seen in the lower fitted value of k. Taken together, also in this condition we can recapitulate experimental observations (see Fig. S10).

**Figure S10:**
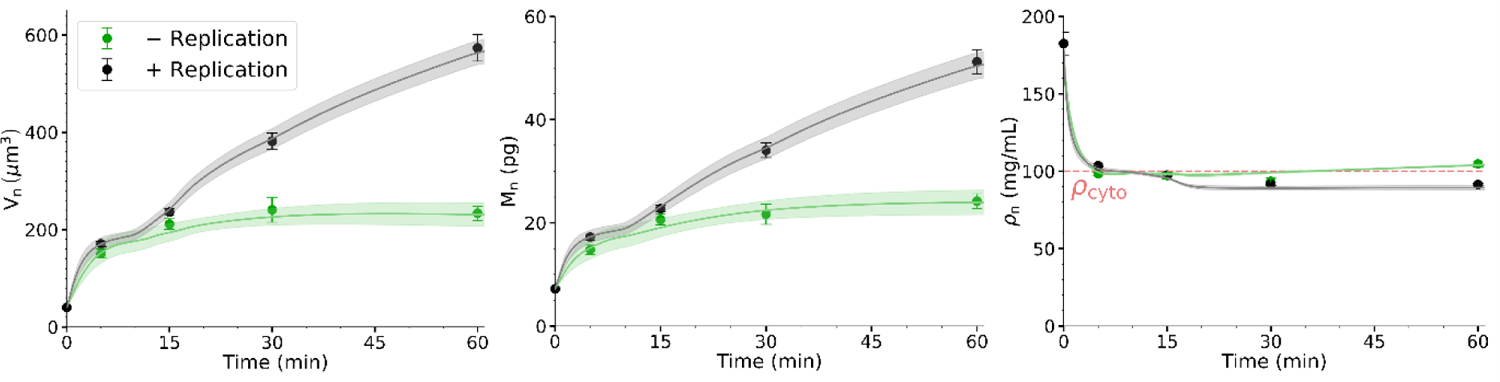
Nuclear volume, dry mass and density as a function of time for nuclei with replication inhibition compared to control condition. Dots represent experimental data and solid lines are the simulation results. Error bars and shaded areas are respectively showing the SEM from experiments and simulations. Respective parameters can be found in Table S2 (-Rep. condition).

The replication inhibition interfered with chromatin via drug treatments. We can, however, change the chromatin content in a more natural way, by using sperm chromatin of a different frog species.

#### 1.4.3. Nuclei assembled around *X tropicalis*sperm chromatin

To evaluate the effect of chromatin content on nuclear volume, dry mass and density, *X. tropicalis (Xt)* sperm, which has a genome half-size of *X laevis (XI),* was used as a chromatin source in the nuclear assembly reaction. Reconstituted *Xt* nuclei were significantly smaller than control X/nuclei (see Fig. S4A). Additionally, these nuclei had a lower dry mass but the same density as control nuclei (see Figs. 4E and G).

One obvious change is to adjust the value of *N_bp_* for the genome of *Xt. A* significantly smaller A? genome results in a lower chromatin pressure, which is essential to reproduce the smaller volumes observed in A?nuclei (see Fig. S4A). Naturally, the value we obtain for *V_chr_* is lower compared to control. The fitted « value for *Xt* is higher compared to control (see Table S2), but considering their errors, we can attribute this difference to natural variations.

Given that the cytoplasmic composition remains the same, we leave the value of *k_in_* unchanged compared to control (see equation (S20)) and only adjust the value of *k’_out_* which depends on chromatin content, in particular the amount of RanGEF which we can expect to be lower compared to *X/*(see Fig. 4F). A lower amount of RanGEF is expected to increase the value of *k’_out_* (see equation (S22)), which leads to a lower value of steady-state nuclear dry mass. Taken together, this choice of parameters leads to the theoretical nuclear density of *Xt* being almost identical to *X/* and a good agreement between experiment and theory (see Fig. S11).

**Figure S11:**
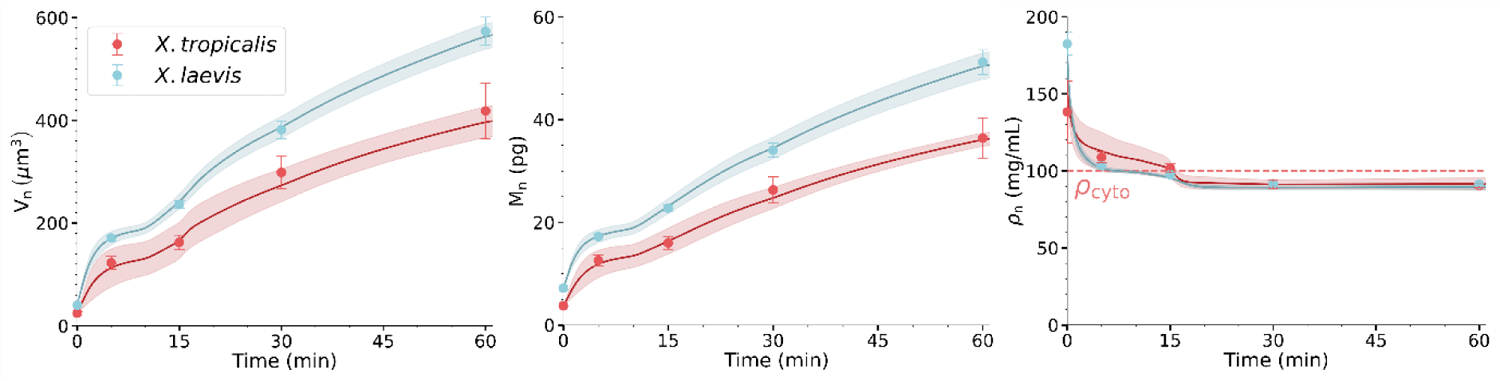
Nuclear volume, dry mass and density as a function of time for nuclei assembled around *X. tropicah’s* sperm chromatin compared to control condition. Dots represent experimental data and solid lines are the simulation results. Error bars and shaded areas are respectively showing the SEM from experiments and simulations. Respective parameters can be found in Table S2 (Trop. condition).

### 1.5. Technical implementation

In this section, we explain in more detail how the theoretical curves shown in the figures of the main text and here were obtained. We start with the parameter *n* that represents chromatin properties. For phase 1, when there is no closed nuclear envelope, we assume the nucleus to be the space occupied by the unfolding chromatin. We describe chromatin as a semi-flexible, self-avoiding polymer. The polymer’s radius of gyration depends on *n* and number of base pairs via *R_g_= N^3/5^* (see 1.1.3). By equating the volume of the sphere with *R = R_g_\o* the measured volume *V_n_* = 4 / 3 R_g_^3^, we thus find the value of *K(t) at time t:*

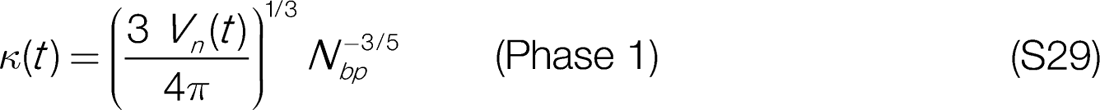

where *V_n_(t)* is the mean nuclear volume measured at time *t*. To find «(/) in phase 2 of nuclear assembly and growth, we numerically solve the pressure balance equation for *k(t)*:

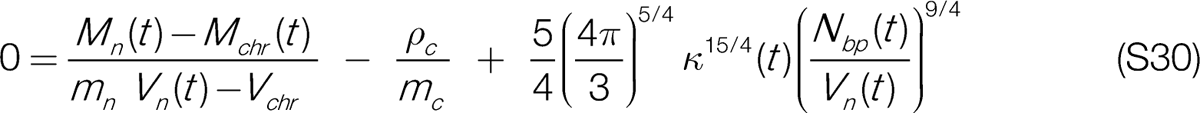

where *V_n_(t)* is the mean nuclear volume measured at time *t, M_n_(t)* the mean nuclear dry mass measured at time *t, M_chr_(t)* the estimated (calculated) chromatin mass at time *t*, and *N_bp_(t)* the number of base pairs of the genome at time *t*. The number of proteins in the nucleus was calculated from the experimental data as: *N_n_(t) = M_n_(t) - M_t_) / m_n_*. Note that *M_chr_(t)* is calculated from equation (S3) and that both *M_chr_(f)* and *N_bp_(t)* double their values after replication.

The times at which the experimental measurements were performed are *t* = 0, 5, 15, 30 and 60 minutes. We define *t* = 10 minutes as the time where phase 2 begins and *k* = k(t*)*. There are limitations to the values that *k\** can take because the corresponding nuclear volume should be smaller than the mean nuclear volume measured at 15 minutes. The upper bound is then k_15_, which is determined by inserting *V_n_(t = 15)* in equation (S29). We then choose:

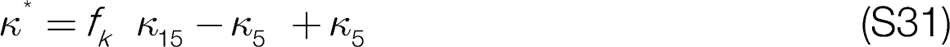

where 0 < *f_k_* < 1 is a simulation parameter that lets us modulate the nuclear volume at the start of phase 2.

We next use *n* values at 0, 5, 15, 30, 60 minutes and at f =10 minutes to find a fit function that describes k(/) at any time point:

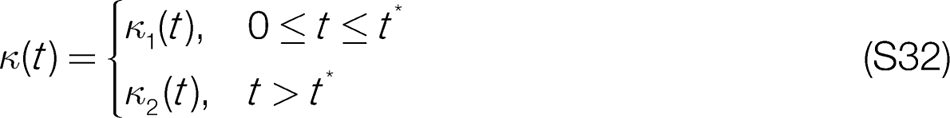

where *K_1_*(t) is a least squares fit function for phase 1 with a Michaelis-Menten saturation curve and *K_2_(t)* for phase 2, with a sigmoidal curve.

Once a function for is obtained, we can use it to solve the system of two ordinary differential equations (ODEs) (S25) and (S26). Given that equation (S26) is non­linear, an analytic solution cannot be obtained and we have to solve these equations numerically. For this purpose, we first bring the system of ODEs to a dimensionless form by choosing the following dimensionless units:

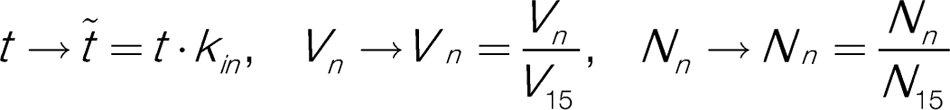

where /V_15_, /_15_ respectively denote the nuclear number of proteins and volume measured at 15 minutes. With these definitions we can bring other quantities to dimensionless form, for instance, for the unit of length *L,* we have *L —»▪ ∼L = L* / l/g^3^. Therefore, the equations that determine the nuclear protein number and volume during phase 1 become:

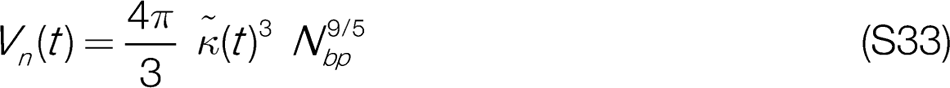

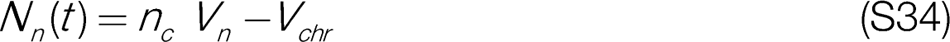

where n_c_ = p_c_ V_15_/m_c_ N_15._ For phase 2, the ODEs have the following dimensionless form:

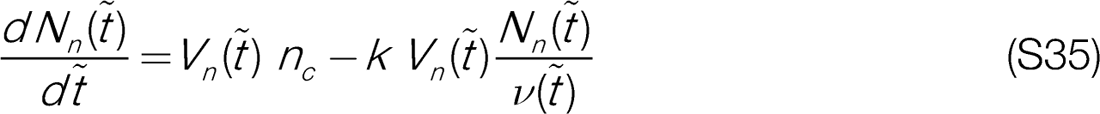

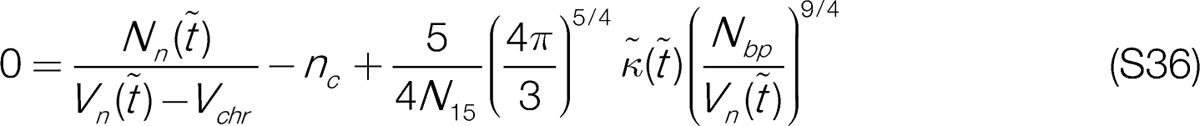

#### 1.5.1. Stochasticity of phase 2 starting time point

Our model includes phases 1 and 2 of nuclear growth, where nuclear volume and protein number are determined by a different set of equations, see equations (S33) and (S34) for phase 1 and (S35) and (S36) for phase 2. We defined a time *f* where the model switches from the phase 1 equations to the phase 2 equations and assumed a value of *t* = 10 minutes based on experimental observations. In reality, the start of phase 2 is somewhat different for each nucleus grown in our system and we included this effect in the following way: for each fitted set of parameters, we performed multiple simulations, where we shifted the phase 2 starting point of f = 10 minutes by a random Gaussian number with mean 0 and a variance of 1 minute. The same shift is applied to the starting and end times of replication. This will of course change the fitted function, but the rest of the parameter values are left unchanged. We perform *N_stoch_* = 50 such stochastic trials and the resulting average curve is shown in the figures for the respective data sets,

#### 1.5.2. Choice of the excluded volume of chromatin

In section 1.1.4, we introduced the parameter *V_chr_,* which represents the chromatin excluded volume in the nucleus. This quantity is in principle time-dependent because the parameter, which is a measure of chromatin condensation, is also time dependent. This would make *V_chr_* another system variable like nuclear volume *V*_n_(t) and protein number *N_n_(t)*. This poses a problem because we only have two equations describing our system, see (S35) and (S36).

For phase 1, we proceed as follows: at *t* = 0, we choose the value of *V_chr_*, for which we obtain the experimentally measured volume and dry mass under the assumption of an osmotic balance. The equation determining *V_chr_* is then:

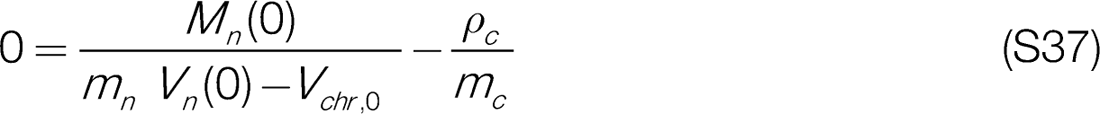

from which it follows:

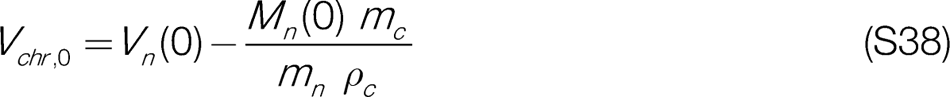

where V_n_(0), *M_n_*(*0*) respectively denote the mean nuclear volume and dry mass measured at *t* = 0. We then change the value of from *V_chr_$* to a different value, which we define as l/_c/r5_, at f = 5 minutes in a linear fashion. For the rest of the simulation, we will assume **V_chr_*(t) =* Therefore, *V_chr_* has the following form:

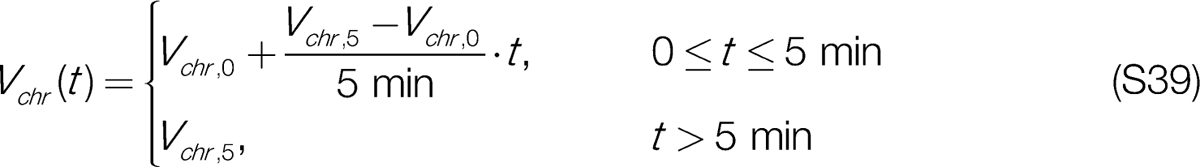

To choose the value of *V_chr5_,* we first calculate *V_chr_* using equation (S38) and the experimental data for *V_n_* and *M_n_* for all time points. This gives us the upper bound estimate for the *V_chr_* under the assumption of zero chromatin pressure. We then set *V_chr,5_* equal to the smallest calculated *V*_chr_. The reason for this is to ensure that nuclear colloid osmotic pressure is less or equal to cytosolic colloid osmotic pressure at all times in the simulations, so that the pressure balance including the contribution of chromatin could be fulfilled. In the majority of cases, the smallest value is indeed just the value calculated for *t* = 5 min, but in some rare cases we have to use the value provided by the formula that corresponds to later times.

#### 1.5.3. Variability in experiments and of simulation parameters

We remarked in section 1.5.1 that the start of phase 2 may vary for nuclei assembled in our experiments. This variability is also present in other important quantities such as the nuclear volume and dry mass. To reproduce this variability also in simulations (and thus gain information on the variability of model parameters), we fitted the theory to each individual experimental data set (for the example shown below, an individual data set is a set of control measurements performed for every perturbation experiment) and then take the average of the obtained theoretical curves, which will also provide us with an error for the theoretical predictions. As we can see in figures S12 and S13 (left panels), the different (control) data sets show significant variability in volume and dry mass, which is reflected in the gray area representing the standard error of mean (right panels) in the theory plot.

**Figure S12:**
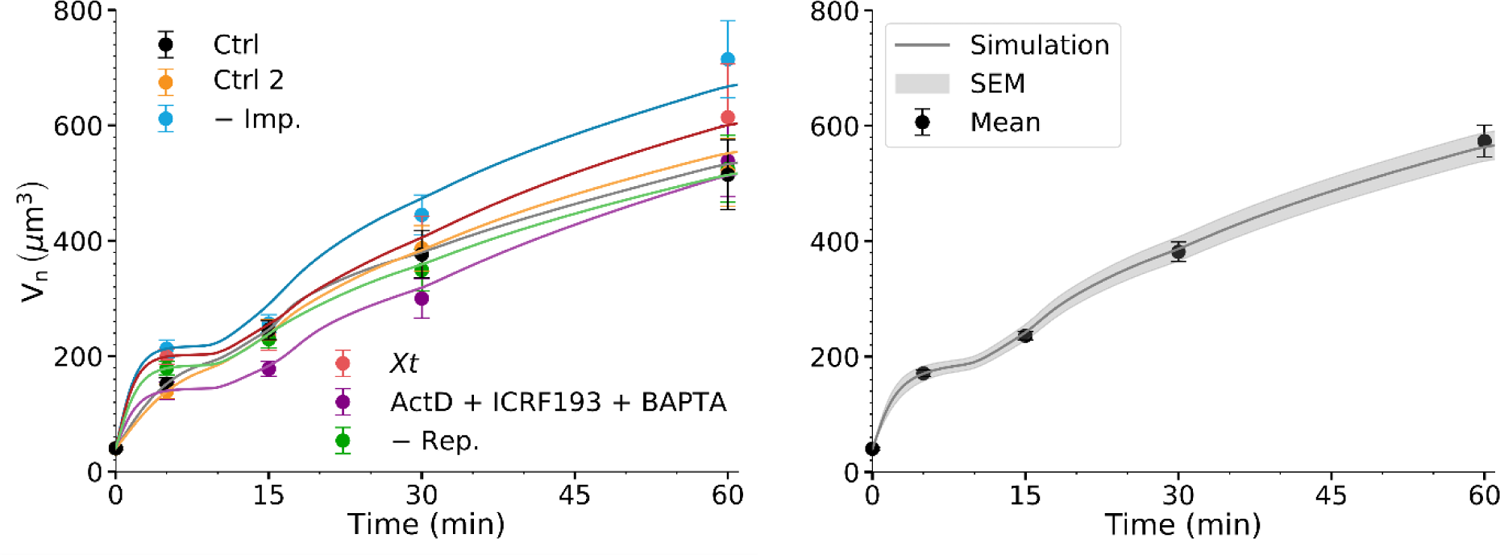
Variability in volume measurements in control conditions and respective theoretical fits. Left panel shows control measurement performed for the respective perturbations (that is why here the perturbation name is used as a label for the respective control set) shown as dots with error bars. Individual theory fits are shown as solid lines. The black data and line labeled as control is the overall average of all control measurements. Right panel shows the average of theoretical fits with shaded region representing SEM.

**Figure S13:**
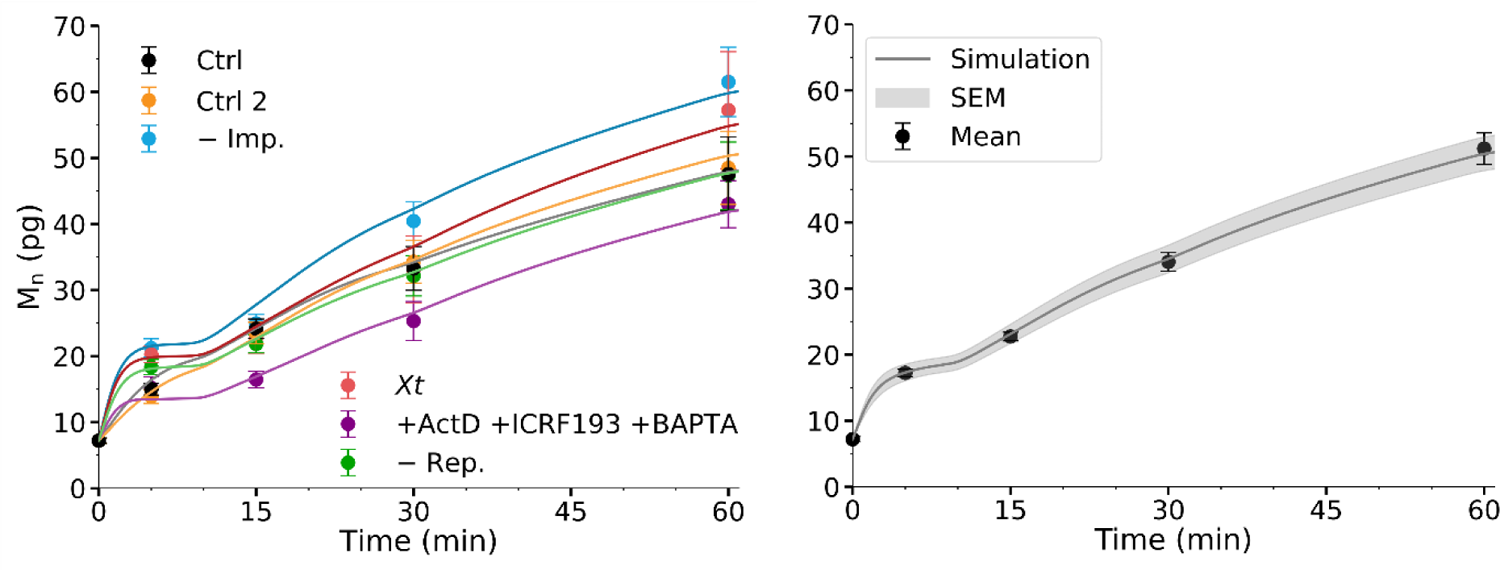
Same as above but for the dry mass

To produce theoretical curves with error bars for the perturbation conditions, we use individual biological replicates as individual data sets. This is how we generated all the plots here and in the main text. For such theoretical fits we can now provide the parameter values with the respective error (SEM) which would indicate variability necessary to capture the natural variability present in the experimental system (see Table S2).

## Footnotes

1 Reality is certainly more complex, with proteins spanning a broad spectrum of nuclear localization degree (from being predominantly cytosolic to exclusively nuclear) and dependent on different transport proteins. Yet, this simplified kinetic approach represents the stereotypic pathway of protein accumulation in the nucleus.

2 Note that nuclear proteins take up only a fraction of cytosolic pool of proteins. Here, we set the cytosolic concentration as the constant concentration of nuclear proteins in the cytosol. The respective proportionality constant, without loss of generality, can be absorbed in the rate constants *k_inout_,* which are parameters in our model.

